# Loss of Intracellular Fibroblast Growth Factor 14 (iFGF14) *Increases* the Excitability of Mature Hippocampal and Cortical Pyramidal Neurons

**DOI:** 10.1101/2024.05.04.592532

**Authors:** Joseph L. Ransdell, Yarimar Carrasquillo, Marie K. Bosch, Rebecca L. Mellor, David M. Ornitz, Jeanne M. Nerbonne

**Affiliations:** Departments of Medicine, Cardiovascular Division, Washington University School of Medicine, St. Louis, MO 63110; Departments of Developmental Biology, Washington University School of Medicine, St. Louis, MO 63110

**Keywords:** iFGF14, Fibroblast growth factor homologous factor 4 (FHF4), Intrinsic excitability, Voltage-gated sodium (Nav) channels, Nav channel activation, Nav channel inactivation

## Abstract

Mutations in *FGF14*, which encodes intracellular fibroblast growth factor 14 (iFGF14), have been linked to spinocerebellar ataxia type 27 (SCA27), a multisystem disorder associated with progressive deficits in motor coordination and cognitive function. Mice (*Fgf14^-/-^*) lacking iFGF14 display similar phenotypes, and we have previously shown that the deficits in motor coordination reflect *reduced* excitability of cerebellar Purkinje neurons, owing to the loss of iFGF14-mediated regulation of the voltage-dependence of inactivation of the fast transient component of the voltage-gated Na^+^ (Nav) current, I_NaT_. Here, we present the results of experiments designed to test the hypothesis that loss of iFGF14 also attenuates the intrinsic excitability of mature hippocampal and cortical pyramidal neurons. Current-clamp recordings from adult mouse hippocampal CA1 pyramidal neurons in acute *in vitro* slices, however, revealed that repetitive firing rates were *higher* in *Fgf14^-/-^*, than in wild type (WT), cells. In addition, the waveforms of individual action potentials were altered in *Fgf14^-/-^* hippocampal CA1 pyramidal neurons, and the loss of iFGF14 reduced the time delay between the initiation of axonal and somal action potentials. Voltage-clamp recordings revealed that the loss of iFGF14 altered the voltage-dependence of activation, but not inactivation, of I_NaT_ in CA1 pyramidal neurons. Similar effects of the loss of iFGF14 on firing properties were evident in current-clamp recordings from layer 5 visual cortical pyramidal neurons. Additional experiments demonstrated that the loss of iFGF14 *does not* alter the distribution of anti-Nav1.6 or anti-ankyrin G immunofluorescence labeling intensity along the axon initial segments (AIS) of mature hippocampal CA1 or layer 5 visual cortical pyramidal neurons *in situ*. Taken together, the results demonstrate that, in contrast with results reported for neonatal (rat) hippocampal pyramidal neurons in dissociated cell culture, the loss of iFGF14 does *not* disrupt AIS architecture or Nav1.6 localization/distribution along the AIS of mature hippocampal (or cortical) pyramidal neurons *in situ*.

## INTRODUCTION

The intracellular fibroblast growth factor subfamily, *Fgf11-14* in mouse and *FGF11-14* in human, generate four iFGF proteins, iFGF11-14, with similar “core” sequences and distinct N-termini (Goldfarb, 2005; Pablo and Pitt, 2016; Ornitz and Itoh, 2022). Alternative exon usage and splicing (Munoz-Sanjuan et a., 2000) generate further iFGF protein diversity. Also referred to as fibroblast growth factor homologous factors, FHF1-4, the iFGFs share sequence and structural homology with the canonical FGFs, but lack the signal sequence for secretion (Smallwood et al, 1996; Hartung et al., 1997; Munoz-Sanjuan et al., 2000; Olsen et al., 2003; Goldfarb, 2005; Pablo and Pitt, 2016; Ornitz and Itoh, 2022). The iFGFs have been shown to bind to the carboxy termini of voltage-gated Na^+^ (Nav) channel pore-forming (α) subunits (Liu et al., 2001, 2003; Wittmack et al., 2004; Lou et al., 2005; Goetz et al., 2009; Wang et al., 2015), and to modulate the time- and voltage-dependent properties of heterologously expressed Nav channels (Liu et al., 2001, 2003; Wittmack et al., 2004; Lou et al., 2005; Rush et al., 2006; Laezza et al., 2007, 2009; Dover et al., 2010; Wang et al., 2011a), as well as of native Nav currents in cardiac (Wang et al., 2011b; Park et al., 2016; Wang et al., 2017; Abrams et al., 2020; Chakouri et al., 2022; Angsutaraux et al., 2023; Fischer et al., 2024) and neuronal (Goldfarb et al., 2007; Dover et al., 2010; Venkatesan et al., 2014; Yan et al., 2014; Bosch et al., 2015; Puranam et al., 2015; Alshammari et al., 2016; Yang et al., 2017; Effraim et al., 2019, 2022) cells.

Unlike canonical FGFs, the iFGFs are broadly and robustly expressed in the adult peripheral and central nervous systems (Smallwood et al., 1996; Wang et al., 2000; Hartung et al., 2007; Ornitz and Itoh, 2022), and mutations in *FGF12 and FGF14* have been linked to inherited disorders of neuronal excitability (van Swieten et al., 2003; Dalski et al., 2005; Brusse et al., 2006; Misceo et al., 2009; Choquet et al., 2015; Al-Mehmadi et al., 2016; Siekierska et al., 2016). *FGF14*, for example, was identified as the locus of mutations in spinocerebellar ataxia type 27, SCA27 (now SCA27A), a rare autosommal dominant neurological disorder which presents with profound and progressive motor and cognitive impairment (van Swieten et al., 2003; Dalski et al., 2005; Brusse et al., 2006; Misceo et al., 2009; Choquet et al., 2015). More recently, GAA repeat expansions in *FGF14* were identified in patients with late onset ataxias, now referred to as SCA27B (Pellerin et al., 2023; Rafehi et al., 2023a,2023b; Mohrens et al., 2024).

Mice (*Fgf14^-/-^*) harboring a targeted disruption of *Fgf14* display profound motor and cognitive deficits (Wang et al., 2002; Xiao et al., 2007; Wozniak et al., 2007), i.e., phenotypes similar to those seen in individuals with SCA27A (Brusse et al., 2006; Misceo et al., 2009). We previously demonstrated that spontaneous, high frequency repetitive firing is markedly attenuated in *Fgf14^-/-^* cerebellar Purkinje neurons (Shakkottai et al., 2009; Bosch et al., 2015), the sole output neurons of the cerebellar cortex (Billard et al., 1993; Gauck and Jaeger, 2000; Ito, 2001). Similar deficits in balance and motor coordination and in spontaneous action potential firing were observed with acute, *in vivo* shRNA-mediated knockdown of *Fgf14* in adult Purkinje neurons (Bosch et al., 2015). Loss of iFGF14, however, did not measurably affect the distribution or intensity of Nav α subunit or Ankyrin G labelling at the axon initial segments (AIS) of mature cerebellar Purkinje neurons. Voltage-clamp experiments revealed that the loss of iFGF14 results in a hyper-polarizing shift the voltage-dependence of inactivation of the transient component of the Nav current (Bosch et al., 2015). Additional experiments revealed that membrane hyperpolarization “rescues” spontaneous repetitive firing in adult cerebellar Purkinje neurons with acute, *Fgf14-*shRNA-mediated knockdown of *Fgf14*, as well as in adult *Fgf14*^-/-^ Purkinje neurons (Bosch et al., 2015). We also showed that viral-mediated expression of iFGF14 in adult *Fgf14*^-/-^ Purkinje neurons rescued spontaneous firing and improved motor coordination and balance (Bosch et al., 2015).

Loss of iFGF14 is also linked to cognitive dysfunction. Individuals with SCA27A, for example, have progressive deficits in learning and memory (Brusse et al., 2006; Misceo et al., 2009) and, compared with wild type (WT) littermates, *Fgf14^-/-^* animals perform poorly in the Morris water maze (Wozniak et al., 2007), suggesting impaired hippocampal functioning. The experiments here were undertaken to test the hypothesis that iFGF14 also modulates Nav current inactivation to regulate the intrinsic excitability of hippocampal pyramidal neurons. We found that evoked repetitive firing rates are *higher* (*not lower*) in *Fgf14^-/-^*, than in WT, hippocampal CA1 pyramidal neurons. In addition, whole-cell voltage-clamp recordings demonstrate that the loss of iFGF14 selectively altered the voltage-dependence of Nav current *activation* (*not inactivation*). In addition, and in contrast with previously published findings on dissociated hippocampal neurons (Pablo et al., 2016), the loss of iFGF14 *does not* disrupt localization of Nav channels at the AIS of mature hippocampal pyramidal neurons *in vivo*.

## MATERIALS AND METHODS

### Animals

All experiments involving animals were performed in accordance with the guidelines published in the National Institutes of Health *Guide for the Care and Use of Laboratory Animals* and all protocols were approved by the Washington University Institutional Animal Care and Use Committee (IACUC). Animals harboring a targeted disruption of the *Fgf14* locus (*Fgf14^-/-^*), described in Wang et al. (2002), were generated by crossing *Fgf14^+/-^* heterozygotes congenic in the C57BL/6J background; genotypes were confirmed by PCR. Adult male and female wild-type (WT) and *Fgf14^-/-^* animals of varying ages (4-10 weeks) were used, as indicated in descriptions of the individual experiments.

### Preparation of Acute Brain Slices

For electrophysiological recordings, acute (350 μM) slices were prepared from (4-8 week old) male and female WT and *Fgf14^-/-^* animals: horizontal sections were prepared for recordings from hippocampal CA1 pyramidal neurons and coronal slices were prepared for recordings from pyramidal neurons in layer 5 of the primary visual cortex, using standard procedures (Davie et al., 2006). Briefly, animals were anaesthetized with 1.25% Avertin and perfused transcardially with ice-cold cutting solution containing (in mM): 240 sucrose, 2.5 KCl, 1.25 NaH_2_PO4, 25 NaHCO_3_. 0.5 CaCl_2_ and 7 MgCl_2_ saturated with 95% O_2_/5% CO_2_. After perfusion, brains were rapidly removed and coronal or horizontal slices were cut in ice-cold cutting solution on a Leica VT1000 S microtome (Leica Microsystems Inc. Buffalo Grove, IL, USA) and incubated in a holding chamber in artificial control cerebrospinal fluid (control ACSF) containing (in mM): 125 NaCl, 2.5 KCl, 1.25 NaH_2_PO_4_, 25 NaHCO_3_, 2 CaCl_2_, 1 MgCl_2_, and 25 dextrose (pH 7.2; ∼310 mOsmol l^−1^), saturated with 95% O_2_/5% CO_2_. Horizontal slices were incubated at 33°C for 25 minutes, followed by at least 35 minutes at room temperature prior to electrophysiological recordings, and coronal slices were incubated for at least 1 hour at room temperature prior to electrophysiological recordings.

### Electrophysiological Recordings

Whole-cell current-clamp recordings were obtained at 33 ± 1°C from visually identified hippocampal CA1 or layer 5 visual cortical pyramidal neurons in acute slices, prepared from adult (6-8 week) WT and *Fgf14^-/-^* animals as described above, using differential interference contrast optics with infrared illumination. Experiments were controlled and data were collected using a Multiclamp 700B patch clamp amplifier interfaced, with a Digidata 1332 acquisition system and the pCLAMP 10 software (Molecular Devices), to a Dell computer. For current-clamp recordings, slices were perfused continuously with warmed (33 ± 1°C) control ACSF, saturated with 95% O_2_/5% CO_2_. Recording pipettes contained (in mM): 120 KMeSO_4_, 20 KCl, 10 HEPES, 0.2 EGTA, 8 NaCl, 4 Mg-ATP, 0.3 Tris-GTP and 14 phosphocreatine (pH 7.25; ∼300 mosmol l^−1^); pipette resistances were 2-4 MΩ. Tip potentials were zeroed before membrane-pipette seals were formed. Pipette capacitances were compensated using the pCLAMP software. Signals were acquired at 50-100 kHz and filtered at 10 kHz prior to digitization and storage. Initial resting membrane potentials (*V*_r_) were between -60 and -80 mV. In each cell, single action potentials and action potential trains were elicited from V_r_ in response to brief (2.5 ms) and prolonged (500 ms), respectively, depolarizing current injections of variable amplitudes.

Voltage-clamp recordings were obtained at 33 ± 1°C from hippocampal CA1 pyramidal neurons in acute slices prepared from young adult (4-6 week) WT and *Fgf14^-/-^* animals, as described above. In this case, recording pipettes (2-4 MΩ) contained (in mM): 110 CsCl, 20 TEA-Cl, 1 CaCl_2_, 2 MgCl_2_, 10 EGTA, 4 Mg-ATP, 0.3 Tris-GTP and 10 HEPES (pH 7.25; ∼310 mosmol/L). After formation of a gigaseal, the control ACSF superfusing the slice was switched to a low Na^+^ ACSF which contained (in mM): 25 NaCl, 100 TEA-Cl, 2.5 KCl, 1.25 NaH_2_PO_4_, 25 NaHCO_3_, 2 CaCl_2_, 1 MgCl_2_, 0.1 CdCl_2_, and 25 dextrose, (pH 7.25; ∼310 mosmol/L), also saturated with 95% O_2_/5% CO_2_. Series resistances were compensated ≥80%, and the voltage errors resulting from the uncompensated resistances were always ≤ 4mV and were not corrected. Voltage-clamp protocols were tested and optimized, using the strategy outlined in Milescu et al. (2010), to minimize space clamp errors and enable reliable measurement of Nav currents in WT and *Fgf14^-/-^* hippocampal CA1 pyramidal neurons.

To determine the voltage-dependence of activation of the transient component of the Nav current, I_NaT_, in WT and *Fgf14^-/-^* CA1 pyramidal neurons, a three-step voltage-clamp protocol (Milescu et al., 2010; Bosch et al., 2015) was used. Briefly, a 5 ms voltage step to 0 mV from a holding potential of -80 mV, to inactivate the Nav currents in the distal neurites, was delivered first, followed by a brief (3 ms) hyperpolarizing voltage step to -60 mV, to allow recovery of the Nav currents in the soma and proximal neurites. Each cell was then depolarized, directly from -60 mV (for 8 ms), to various test potentials to evoke the Nav currents. The voltage-clamp paradigm and a representative set of records, with selected voltage steps shown and color-coded with the corresponding current traces, are presented in **Figure 4A**. The peak inward Nav currents evoked at each test potential in each cell were measured; Nav conductances (G_Na_) at each test potential were determined (using the calculated Na^+^ reversal potential of + 38 mV) and normalized to the maximal Nav conductance (G_Na,max_). Mean ± SEM normalized Nav conductances were then plotted as a function of the test potential and fitted with single (equation 1) or double (equation 2) Boltzmanns using the following equations:

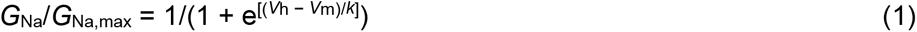

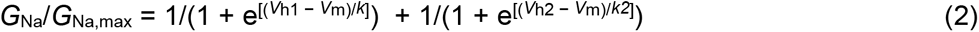

Where *V*_h_ is the potential of half-maximal activation, V_m_ is the test potential, and *k* is the slope factor.

To determine the voltage-dependence of steady-state inactivation of I_NaT_ in *Fgf14^-/-^*and WT CA1 pyramidal neurons, a three step voltage-clamp protocol (Milescu, 2010; Bosch et al., 2015) was also used. In this case, each cell was first depolarized to 0 mV for 5 ms to inactivate the Nav currents, subsequently hyperpolarized briefly (3 ms) to various conditioning potentials to allow recovery and, finally, depolarized to 0 mV for 5 ms. The voltage-clamp paradigm and a representative set of records, with selected voltage steps shown and color-coded with the corresponding current traces, are presented in **Figure 4B**. The peak transient Nav current amplitudes, evoked during the final voltage step to 0 mV from each conditioning potential in each cell were measured and normalized to the current amplitude evoked from the -130 mV conditioning voltage step (in the same cell). Mean ± SEM normalized Nav current amplitudes were then plotted as a function of the conditioning voltage and fitted with the Boltzmann equation:

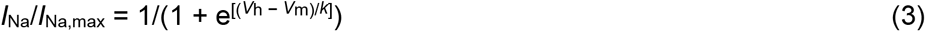

Where *V*_h_ is the potential of half-maximal inactivation, V_m_ is the conditioning potential, and *k* is the slope factor.

### Analyses of Electrophysiological Data

Whole-cell current- and voltage-clamp data were analyzed using ClampFit (Molecular Devices), Microsoft Excel, Mini Analysis (v. 6.0, Synaptosoft, Inc., Decatur, GA, USA) and Prism (GraphPad Software Inc., La Jolla, CA, USA). In all experiments, input resistances (*R*_in_) were determined from the change in membrane potential produced by a 20 pA hyperpolarizing current injection from V_r_. The current threshold for action potential generation in each cell was defined as the minimal current injection, applied (for 2.5 ms) from V_r_, required to evoke a single action potential. The voltage threshold for action potential generation was determined as the voltage at which the action potential slope (dV/dt) was ≥30 mV/ms. Action potential durations at half-maximal amplitude (APD_50_) were determined from measurements of the widths of action potentials when the membrane voltage returned halfway to V_r_ (from the peak).

Results obtained in the analyses of the waveforms of action potentials and the repetitive firing properties of individual hippocampal CA1 and layer 5 visual cortical pyramidal neurons are displayed, and averaged/normalized data are presented as means ± SEM. Statistical analyses were performed using a Student’s (unpaired) *t* test unless otherwise noted and (exact) *P* values are presented in the text, in Table 1 and/or the Figure legends.

**Table 1.** Resting and active membrane properties of WT and *Fgf14^-/-^* hippocampal CA1 and layer 5 cortical pyramidal neurons^1^.

| Cell Type | R<sub>in</sub> (MΩ) | V<sub>rest</sub> (mV) | AP<sub>Thr</sub> (mV) | APA (mV) | APD<sub>50</sub> (ms) | dV<sub>2</sub>/dt interval (ms) | Peak +dV/dt (mV/ms) |
| --- | --- | --- | --- | --- | --- | --- | --- |
| WT CA1<br>(n = 13) | 62 ± 7 | -67 ± 2 | -50 ± 1 | 106 ± 3 | 0.84 ± .03 | 0.16 ± .01 | 459 ± 23 |
| *Fgf14*<sup>-/-</sup> CA1<br>(n = 14) | 69 ± 4 | -68 ± 1 | -46 ± 1 | 107 ± 2 | 0.71 ± .02 | 0.09 ± .01 | 573 ± 29 |
| *P* value | 0.382 | 0.291 | 0.015 | 0.924 | < 0.001 | < 0.001 | 0.007 |
| WT Cortex<br>(n = 17) | 76 ± 8 | -68 ± 1 | -44 ± 1 | 97 ± 2 | 0.67 ± .03 | 0.08 ± .01 | 458 ± 28 |
| *Fgf14*<sup>-/-</sup> Cortex<br>(n = 12) | 66 ± 8 | -69 ± 1 | -49 ± 1 | 103 ± 2 | 0.53 ± .02 | 0.07 ± .01 | 624 ± 31 |
| *P* value | 0.342 | 0.620 | 0.009 | 0.342 | 0.009 | 0.231 | < 0.001 |

<sup>1</sup>All data are means ± SEM; n = number of cells. R<sub>in</sub> = input resistance; V<sub>rest</sub> = resting membrane potential; AP<sub>Thr</sub> = voltage threshold for action potential generation; APA = action potential amplitude; APD<sub>50</sub> = action potential duration at 50% repolarization; dV<sub>2</sub>/dt interval = time between the first and second peaks in the second derivative of the voltage (dV<sub>2</sub>/dt) plot; Peak +dV/dt = maximum rate of change of the membrane voltage during the rising phase of the action potential.

### Immunostaining

Using previously described methods (Bosch et al., 2015), adult (8-10 week) WT and *Fgf14^-/-^* mice were anesthetized with 1 ml/kg (intra peritoneal injection) ketamine/xylazine and perfused transcardially with 0.9% NaCl, followed by freshly prepared, ice-cold 1% formaldehyde in 0.1M phosphate buffered saline (PBS) at pH 7.4. Brains were removed and post-fixed (1% formaldehyde in 0.1M PBS) for 1 hr at 4°C. Following washing with PBS, brains were cryoprotected in 30% sucrose in PBS overnight at 4°C, subsequently embedded in Tissue Tek OCT (Sakurai Finetek), and frozen. Horizontal (40 μm) cryostat sections of the hippocampus or cortex were cut, collected in PBS at room temperature (22– 23°C) and stored at 4°C until processed for immunostaining.

Free-floating sections were rinsed twice in PBS, followed by 30 min in PBS with 0.1% Triton X-100 (v/v). Sections were then incubated in blocking buffer (PBS plus 10% goat serum) for 1 hour, followed by overnight incubation at 4 °C with primary antibodies, diluted (1:1000) in PBS with 0.1% Triton X-100 and 0.1% bovine serum albumin. The primary antibodies used were purified mouse monoclonal Neuromab antibodies obtained from Antibodies, Inc. (Davis, CA): anti-iFGF14 (mAb-α-iFGF14; clone N56/21, IgG1); anti-Ankyrin G (mAb-α-Ankyrin G; clone N106/36, IgG2a), and anti-Nav1.6 (mAb-α- Nav1.6; clone K87A/10, IgG1). Following washing in PBS, sections were incubated with goat anti-mouse secondary antibodies conjugated to Alexa Fluor 488, Alexa Fluor 594 or Alexa Fluor 647 (1:400; Life Technologies) in PBS for one hour at room temperature. Sections were washed in PBS, mounted on positively charged slides, coated with a drop of Vectashield Hardset (Vector Laboratories) and cover slipped; slides were stored at 4°C.

### Image Acquisition and Analysis

Images were captured using an Olympus Fluoview-500 confocal microscope with a 60X oil immersion objective. Laser intensity, gain and pinhole size were kept constant across experiments and data were acquired and analyzed blinded to experimental group. Sequential acquisition of data from multiple channels was used and z-stacks were collected in 0.5 μm steps and converted into maximum intensity *z*-projections using NIH ImageJ. Line scan graphs were also prepared using ImageJ, and integrated intensity plots were generated in MATLAB using previously described methods (Grubb and Burrone, 2010; Bosch et al., 2015). With ImageJ, three channel TIFF files were analyzed from line profiles drawn along the axon initial segment (AIS) beginning at the soma and using the anti-Ankyrin G staining to indicate (mark) the AIS. Fluorescence intensity plot profiles were obtained from each channel and data were imported into Excel for normalization and averaging. The start and end of each AIS was determined by the proximal and distal AIS points, respectively, at which the normalized and smoothed anti-Ankyrin G fluorescence intensity declined to 0.33 of the maximal value (in the same AIS). Fluorescence intensities, obtained from these line profiles (Grubb and Burrone, 2010; Bosch et al., 2015), were smoothed using a 40-point sliding mean. Integrated fluorescence intensities along each AIS were calculated as the area under the curve from the beginning to the end of the AIS, and normalized to the maximal and minimal values in the same AIS; mean ±SEM normalized values are reported.

### Quantitative RT-PCR

For RNA isolation, hippocampal CA1 and primary visual cortical tissue pieces were dissected from anesthetized (1.5% avertin) adult (8-10 week) female and male *Fgf14^-/-^* and WT animals and immediately flash-frozen (individually) in liquid nitrogen. Total RNA was isolated from each tissue sample using Trizol and DNase-treated using the RNeasy Tissue Mini Kit (Qiagen) according to the manufacturer’s instructions. The expression of transcripts encoding iFGF11-14 was determined by SYBR green quantitative RT-PCR using sequence specific primers (**Supplemental Table 1**). Data were analyzed using the threshold cycle (C_T_) relative quantification (2^-ΔCt^) method (Livak and Schmittgen, 2001; Marionneau et al., 2008) using hypoxanthine guanine phosphoribosyl transferase I (hprt) as the endogenous control.

### Western Blot Analysis

Lysates were prepared from frozen hippocampal CA1 and primary visual cortical tissues, obtained as described above from adult (8-10 week) WT and *Fgf14^-/-^* animals, in 20 mM HEPES + 150 mM NaCl buffer with 0.5% CHAPS and a protease inhibitor tablet (Roche), using established methods (Brunet et al., 2004, Norris and Nerbonne, 2010). Immunoprecipitations (IPs) were also performed from the same lysates with the (Neuromab) antibody mAb-α-iFGF14 conjugated to protein G dynabeads (Life Technologies), using previously described methods (Bosch et al., 2016). The cortical/hippocampal lysates and the proteins eluted from the beads following the IPs were fractionated on SDS-PAGE (12%) gels, transferred to polyvinylidene fluoride (PVDF) membranes (Biorad) and probed for iFGF14 expression using a rabbit polyclonal anti-iFGF14 antiserum (Rb-α-iFGF14; 1:1,000; Covance Laboratories), validated as described previously (Bosch et al., 2016). Additional lysates were prepared from frozen hippocampal and primary visual cortical tissues from adult (8-10 week) WT and *Fgf14^-/-^* animals, fractionated, transferred, and probed for iFGF12 and iFGF13 expression using validated (Angsutararux et al., 2023; Nerbonne, unpublished) rabbit polyclonal anti-iFGF12 (Rb-α-iFGF12; 1:500; Sigma) and anti-FGF13 (Rb-α-iFGF13; 1:1000; generous gift of Geoffrey Pitt, Weill Medical College, Cornell University) antibodies. To ensure equal loading of all lanes of the gels in these latter experiments, membranes were also probed with a mouse monoclonal antibody targeting glyceraldehyde-3-phosphate dehydrogenase (mAb-α-GAPDH; 1:8000; ThermoFisher).

## RESULTS

### Robust expression of *Fgf14*/iFGF14 in adult mouse hippocampus and cortex

In addition to expression in the cerebellum, previous *in situ* hybridization studies identified *Fgf14*/ *FGF14* transcripts in adult mouse and human hippocampus and cerebral cortex, as well as in the amygdala, striatum, and thalamus (Wang et al., 2000, 2002). In initial experiments here, we directly examined *Fgf14* transcript and iFGF14 protein expression in the CA1 region of the hippocampus and in the primary visual cortex of adult WT mice. As illustrated in **Figure 1**, quantitative RT-PCR analyses confirmed expression of *Fgf14* transcripts in the CA1 region of the hippocampus (**Figure 1A**) and in the primary visual cortex (**Figure 1B**) of WT animals, and the complete loss of *Fgf14* in *Fgf14^-/-^* hippocampus (**Figure 1A**) and cortex (**Figure 1B**). Also, consistent with previous findings (Wang et al., 2002), *Fgf14B* is the predominant splice variant in both WT tissues, and *Fgf14A* is barely detectable (**Figure 1A** and **1B**). Western blot analyses, using a polyclonal anti-iFGF14 antiserum (Bosch et al., 2015, 2016), confirmed robust expression of the iFGF14 protein in the CA1 region of hippocampus (**Figure 1A**) and in the primary visual cortex (**Figure 1B**) of adult WT mice and the complete loss of the iFGF14 protein in the tissues from *Fgf14^-/-^* animals. The full western blot image is presented in **Supplemental Figure 1**. Additional RT-PCR and Western blot analyses revealed robust expression of *Fgf12* (specifically *Fgf12B*) and *Fgf13* (specifically *Fgf13A*) transcripts and the iFGF12 and iFGF13 proteins in the CA1 region of the hippocampus (**Supplemental Figure 2**) and in the primary visual cortex (**Supplemental Figure 4**) of adult WT mice. Interestingly, in spite of the complete loss of *Fgf14* transcripts and the iFGF14 protein (**Figure 1**), no remodeling (upregulation) of the *Fgf12*/*Fgf13* transcripts and/or the iFGF12/iFGF13 proteins is evident in the CA1 region of the hippocampus (**Supplemental Figure 2**) or the primary visual cortex (**Supplemental Figure 4**) of adult *Fgf14^-/-^* animals (see **Discussion**).

**Figure 1.**
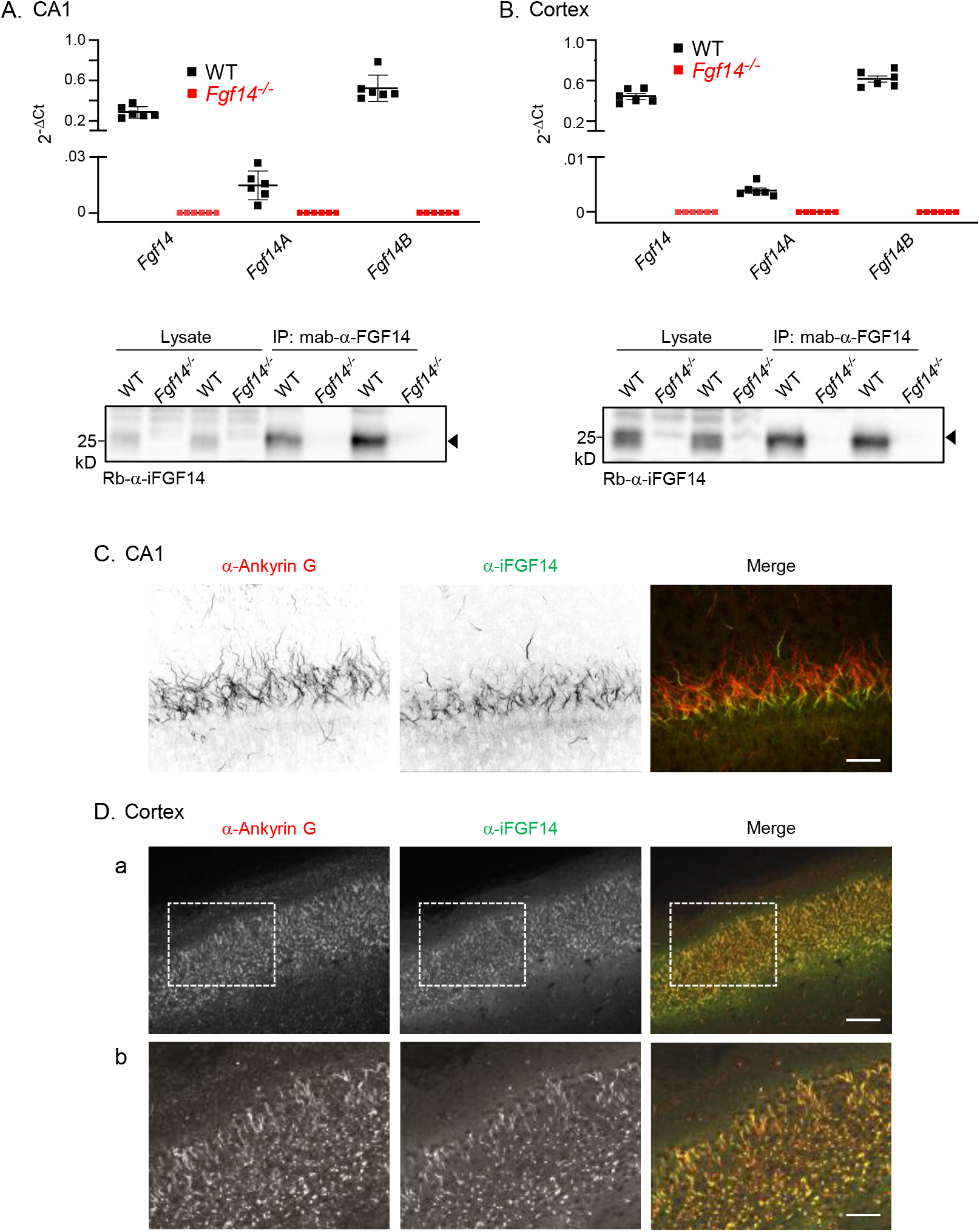
Robust expression of *Fgf14* and iFGF14 in adult mouse hippocampus and cortex. (**A, B**) *Top panels*: qRT-PCR analyses of *Fgf14* transcript expression in the CA1 region of the hippocampus (**A**) and primary visual cortex (**B**) of adult WT (●; n = 6) and *Fgf14^-/-^* (▪, n = 6) mice demonstrate robust expression of *Fgf14* in WT tissues and confirm that *Fgf14* is undetectable in *Fgf14^-/-^* tissues. Further analysis, using isoform specific primers (see: **Supplemental Table 1**), revealed ∼ 60 (**A**) to 100 (**B**) higher expression of the *Fgf14B* transcript, compared with the *Fgf14A* transcript, in the CA1 region of the hippocampus (**A**) and the primary visual cortex (**B**). *Bottom panels*: Western blots of fractionated total protein lysates and proteins immunoprecipitated with a mouse monoclonal anti-FGF14 antibody (mAb-α- FGF14; Neuromab) from the CA1 region of the hippocampus (**A**) and the primary visual cortex (**B**) of adult WT and *Fgf14^-/-^* mice, probed with a rabbit polyclonal anti-iFGF14 antiserum (Rb-α-FGF14), are shown. The arrowheads indicate the iFGF14 protein (missing in the samples from the *Fgf14^-/-^*brains). Full Western blots are presented in **Supplemental Figure 1**. (**C,D**) Representative images of anti-ankyrin G and anti-iFGF14 immunolabeling in the CA1 region of the hippocampus (**C**) and in the primary visual cortex (**D**) of adult WT mice, probed with monoclonal anti-Ankyrin G (mAb-α-Ankyrin G) and anti-iFGF14 antibodies, demonstrate that anti-iFGF14 labelling is localized with anti-Ankyrin G at the axon initial segments of hippocampal CA1 neurons (**C**) and visual cortical neurons (**D**). Dashed boxes in (**Da**) are displayed at a higher magnification in (**Db**); scale bars are 25 μm in (**C**), 50 µm in (**Da**) and 25 μm in (**Db**).

Immunohistochemical experiments with a monoclonal anti-iFGF14 antibody (Xiao et al., 2013; Bosch et al., 2015) revealed anti-iFGF14 labeling at the AIS of WT hippocampal CA1 pyramidal neurons (**Figure 1C**), identified using an anti-Ankyrin G antibody (Bosch et al., 2015). Similar patterns of anti-FGF14 and anti-Ankyrin G labeling were observed in pyramidal neurons in WT mouse primary visual cortex (**Figure 1D**). Confirming robust expression of iFGF14 in adult mouse hippocampal and cortical pyramidal neurons, subsequent experiments were focused on testing the hypothesis that, similar to our findings in cerebellar Purkinje neurons (Bosch et al., 2015), loss of iFGF14 attenuates the intrinsic excitability of mature hippocampal and cortical pyramidal neurons.

### Loss of iFGF14 *increases* evoked repetitive firing rates in hippocampal CA1 pyramidal neurons

To determine the functional effects of the loss of iFGF14 on the excitability of adult CA1 hippocampal pyramidal neurons, whole-cell current-clamp recordings were obtained from visually identified CA1 pyramidal neurons in acute slices prepared from adult (6-8 week) WT and *Fgf14^-/-^*mice, as described in **Materials and Methods.** As illustrated in the representative recordings shown in **Figure 2A** (left panel), WT CA1 pyramidal neurons fired repetitively in response to prolonged (500 ms) depolarizing current injections, and the rates of repetitive firing increased as a function of the amplitude of the injected current. Similar to WT cells, prolonged depolarizing current injections elicited repetitive firing in *Fgf14^-/-^*CA1 pyramidal neurons (**Figure 2A**, right panel), and firing rates also tracked the amplitudes of the injected currents. As is also evident in the representative records shown in **Figure 2A**, repetitive firing rates were higher in the *Fgf14^-/-^*, compared with the WT, CA1 pyramidal neuron for all injected current amplitudes. Similar results were obtained in recordings from additional WT and *Fgf14^-/-^* CA1 pyramidal neurons, and the mean ± SEM input-output curves (numbers of spikes evoked plotted versus injected current amplitudes) in WT (n = 13) and *Fgf14^-/-^* (n = 14) hippocampal CA1 pyramidal neurons (**Figure 2B**) are distinct (*P* = 1 x 10^-6^; two-way ANOVA).

**Figure 2.**
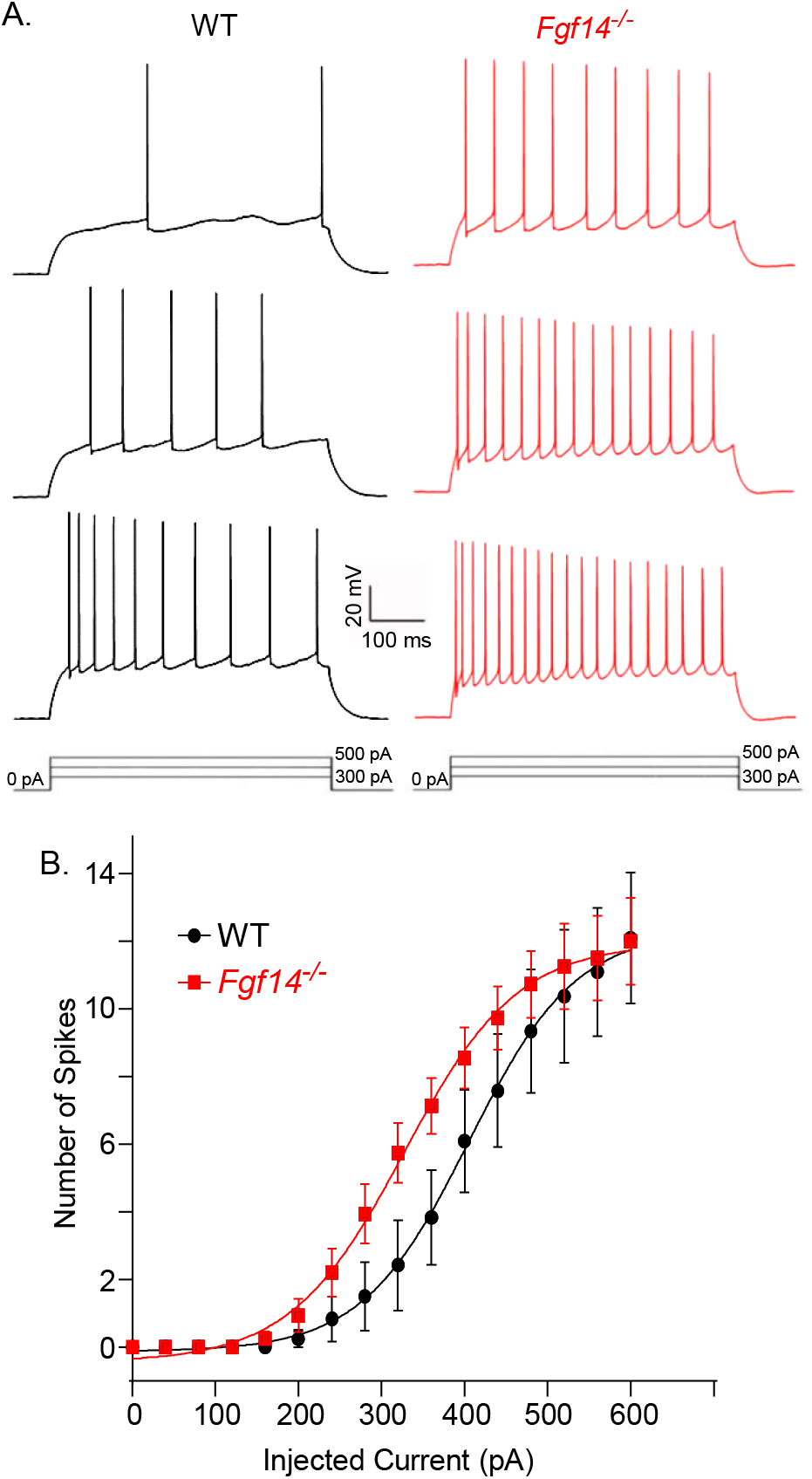
Loss of iFGF14 increases the repetitive firing rates of adult hippocampal CA1 pyramidal neurons. Repetitive firing was evoked in hippocampal CA1 pyramidal neurons in acute slices, prepared from adult (6-8 week) WT and *Fgf14^-/-^* animals, in response to prolonged (500 ms) depolarizing current injections of varying amplitudes (see: **Materials and Methods**). Representative voltage recordings are presented in (**A**) with the injected current amplitudes shown below the voltage records. (**B**) Mean ± SEM numbers of action potentials evoked during 500 ms current injections are plotted as a function of the amplitudes of the injected currents. The resulting input-output curves (**B**) for *Fgf14^-/-^* (▪; n = 15) and WT (●; n = 13) CA1 pyramidal neurons are distinct: *Fgf14^-/-^* (▪) CA1 pyramidal neurons fire at higher (*P* = 1 x 10^-6^; two-way ANOVA) rates than WT (●) CA1 pyramidal neurons at all injected current amplitudes.

Although spike frequency adaptation is apparent in the representative recordings from both WT and *Fgf14^-/-^* neurons (**Figure 2A**), quantification of ratio of the average firing frequency at the end (last 100 ms) and at the beginning (first 100 ms) for current injections that evoked the same mean number of action potentials (400 pA in *Fgf14^-/-^* cells and 460 pA in WT cells, see Figure 2B) revealed similar mean ± SEM firing rate ratios of 0.44 ± .07 (n = 10) and 0.52 ± 0.05 (n = 10) in *Fgf14^-/-^* and WT neurons, respectively. Loss of iFGF14, therefore, does not measurably affect spike frequency adaptation in mature hippocampal CA1 pyramidal neurons.

### Waveforms of individual action potentials are altered in *Fgf14^-/-^* CA1 pyramidal neurons

Additional experiments and analyses were completed to quantify the effects of the loss of iFGF14 on the waveforms of individual action potentials in CA1 pyramidal neurons. As illustrated in **Figure 3A**, the waveforms of single action potentials evoked in *Fgf14^-/-^* and WT CA1 pyramidal neurons in response to brief (2.5 ms) depolarizing current injections are similar (**Figure 3A**). Analyses of the voltage records revealed, however, that there are differences in the voltage thresholds (V_thr_) for action potential generation (**Figure 3B**) and in action potential durations (APD_50_) at 50 % repolarization (**Figure 3C**) in *Fgf14^-/-^* and WT CA1 pyramidal neurons. The mean ± SEM V_thr_ for action potential generation determined in *Fgf14^-/-^* CA1 pyramidal neurons, for example, is depolarized relative to the value measured in WT CA1 cells (**Figure 3B**; **Table 1**). In addition, the mean ± SEM APD_50_ value is smaller in *Fgf14^-/-^*, than in WT, CA1 pyramidal neurons (**Figure 3C**; **Table 1**).

**Figure 3.**
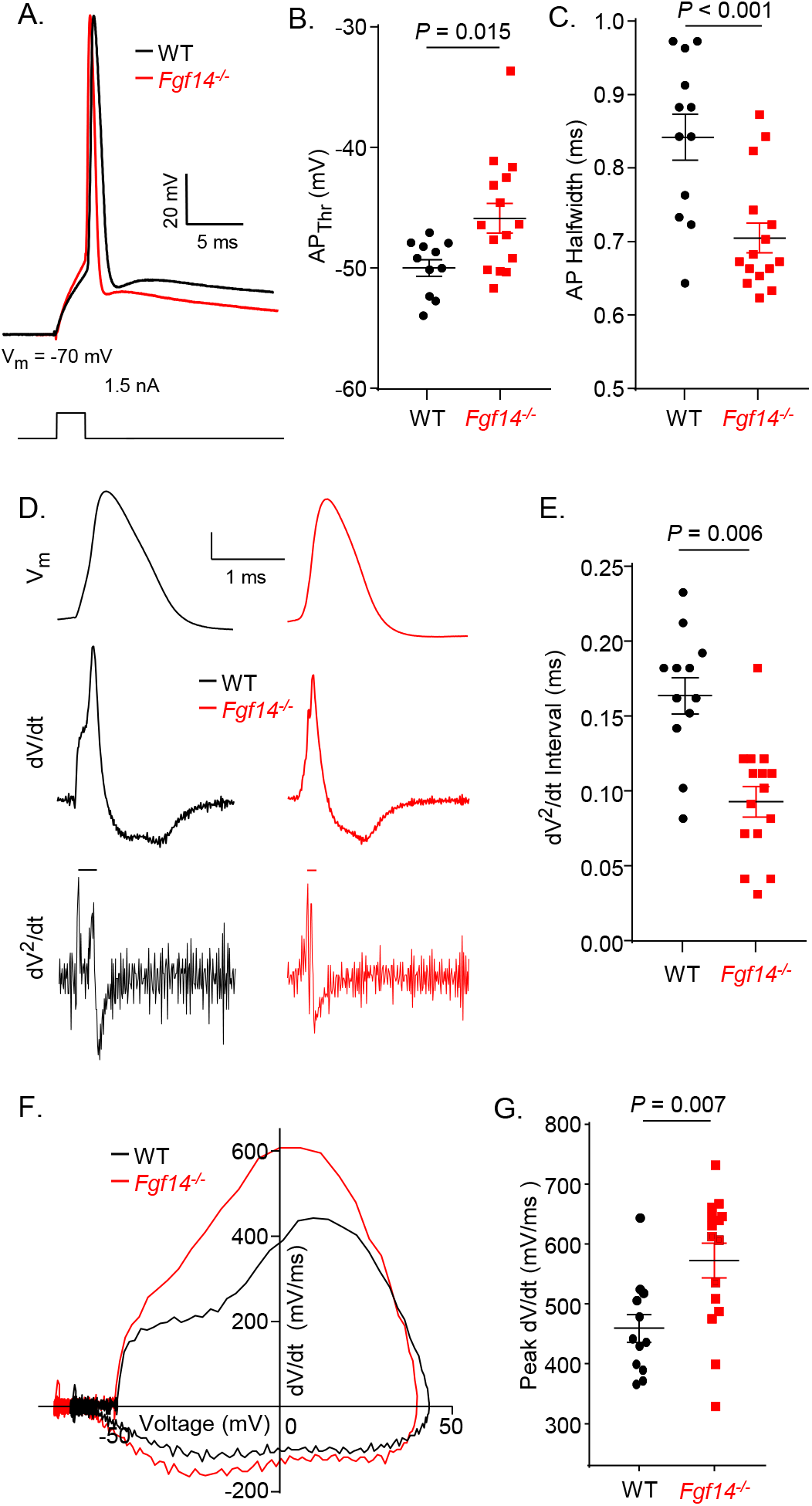
Waveforms of individual action potentials in adult *Fgf14^-/^* and WT hippocampal CA1 pyramidal neurons are distinct. (**A**) Representative recordings of individual action potentials, evoked by brief (2 ms) depolarizing current injections, in WT (*black*) and *Fgf14^-/-^* (*red*) CA1 pyramidal neurons in acute hippocampal slices, prepared from adult (6-8) week animals, are shown. Similar results were obtained in recordings from many CA1 pyramidal neurons and the mean ± SEM voltage threshold for action potential generation (AP_Thr_) was more depolarized (**B**) and the mean ± SEM action potential duration at halfwidth (AP_Halfwidth_) (**C**) was shorter (see: **Table 1**) in *Fgf14^-/-^* (▪; n = 14), than in WT (●; n = 13), cells. (**D**) The representative action potentials in (**A**) are shown on an expanded time scale (*top panels*), and the first (dV/dt; *middle panels*) and second (dV^2^/dt; *lower panels*) derivatives of the action potentials are illustrated in the panels immediately below. The voltage trajectories (dV/dt) of individual action potentials (*middle panels*) are similar in WT and *Fgf14^-/-^* CA1 pyramidal neurons, although a clear difference is evident in the rising phases. In the second derivatives (dV^2^/dt) of the voltage trajectories (*lower panels*), two inflection points, corresponding to the axonal and somal spikes (see text), are clearly evident in both WT and *Fgf14^-/-^* hippocampal CA1 pyramidal neurons, although the temporal relation between them (in *Fgf14^-/-^* and WT cells) is different. As illustrated in (**E**), similar results were obtained in many cells, and the mean interval between the inflection points in the dV^2^/dt versus time plots in *Fgf14^-/-^*(▪; n = 14) CA1 pyramidal neurons is much smaller (see: **Table 1**) than in WT (●; n = 13) CA1 pyramidal neurons. (**F**) Construction of dV/dt versus voltage (phase) plots also revealed clear differences in the voltage dependences of the rising phases of the action potentials in WT and *Fgf14^-/-^* CA1 pyramidal neurons. As illustrated in (**G**), similar results were obtained in many cells, and the mean ± SEM peak dV/dt value for action potentials is higher (see: **Table 1**) in *Fgf14^-/-^*(▪; n = 14), than in WT (●; n = 13), CA1 pyramidal neurons.

Although the membrane voltage (V_m_) trajectories during individual action potentials are similar in WT and *Fgf14^-/-^* CA1 pyramidal neurons (**Figure 3D**, top traces), there are marked differences in the first (dV/dt) (**Figure 3D**, middle traces) and second (dV^2^/dt) (**Figure 3D**, bottom traces) derivative plots of action potential trajectories versus time. In the representative WT CA1 pyramidal neuron presented in **Figure 3D**, for example, there are two distinct components in the rising phase of the dV/dt plot (**Figure 3D**, black middle trace), and these two inflection points, which correspond to action potentials generated in the axon and the cell soma (Palmer and Stuart, 2006; Meeks and Mennerick, 2007), are clearly resolved in the dV^2^/dt plot (**Figure 3D**, black lower trace). Although also evident in the dV/dt (**Figure 3D**, red middle trace) and dV^2^/dt (**Figure 3D**, red lower trace) plots generated from the action potential recorded from a representative *Fgf14^-/-^* CA1 pyramidal neuron (**Figure 3A**), the time difference between the two inflection points, is much smaller in the *Fgf14^-/-^*, than in the WT, cell (**Figure 3D**). Similar results, obtained in recordings from additional WT and *Fgf14^-/-^* CA1 pyramidal neurons, are plotted in **Figure 3E**. As illustrated, the mean ± SEM time interval between the two inflection points is much smaller (*P* = 0.006) in *Fgf14^-/-^*, than in WT, hippocampal CA1 pyramidal neurons (**Figure 3E**, **Table 1**). Phase plots (**Figure 3F**) also revealed that peak dV/dt values are larger (*P* = 0.007) in *Fgf14^-/-^*, compared with WT, CA1 pyramidal neurons (**Figure 3G**; **Table 1**).

### Loss of iFGF14 selectively affects Nav current activation in CA1 pyramidal neurons

To explore the contribution of iFGF14-mediated effects on voltage-gated sodium (Nav) currents to the observed differences in the waveforms of individual action potentials (**Figure 3**) and the repetitive firing rates (**Figure 2**) of *Fgf14^-/-^* and WT CA1 pyramidal neurons, whole-cell Nav currents were recorded from cells in acute slices prepared from young adult (4-6 week) animals (see: **Materials and Methods**). Voltage-clamp experiments were performed using a previously described experimental strategy (Milescu et al., 2010), optimized to minimize space clamp errors and to allow reliable measurements of the voltage-dependences of activation and steady-state inactivation of the fast, transient Nav currents, I_NaT_. To compare the voltage-dependences of activation of I_NaT_ in *Fgf14^-/-^* and WT CA1 pyramidal neurons, mean (± SEM) normalized Nav conductances (G_Na_) were calculated from the measured Nav currents (**Figure 4A**), plotted as a function of the test potential, and fitted with Boltzmann functions (**Figure 4C**). As is evident in **Figure 4C**, the normalized peak I_NaT_ conductance data in WT CA1 pyramidal neurons (n = 8) was best described by a double Boltzmann function (1^st^ term V_1/2_ = -49.0 ± 1.0 mV, 1^st^ term slope factor = 0.8 ± 0.1, 2^nd^ term V_1/2_ = -29.2 ± 1.4 mV, 2^nd^ term slope factor = 6.8 ± 1.1), suggesting that there are two populations of I_NaT_ channels with distinct voltage-dependences of activation in WT CA1 pyramidal cells. The mean ± SEM normalized conductance data for I_NaT_ in *Fgf14^-/-^* CA1 pyramidal neurons (n = 8), in contrast, was well-described by a single Boltzmann with a V_1/2_ = -39.1 ± 1.0 mV and a slope factor = 7.1 ± 1.0 (**Figure 4C**), values that are distinct from those observed in WT cells, suggesting that both populations of Nav channels in WT CA1 pyramidal neurons are affected by the loss of the iFGF14.

**Figure 4.**
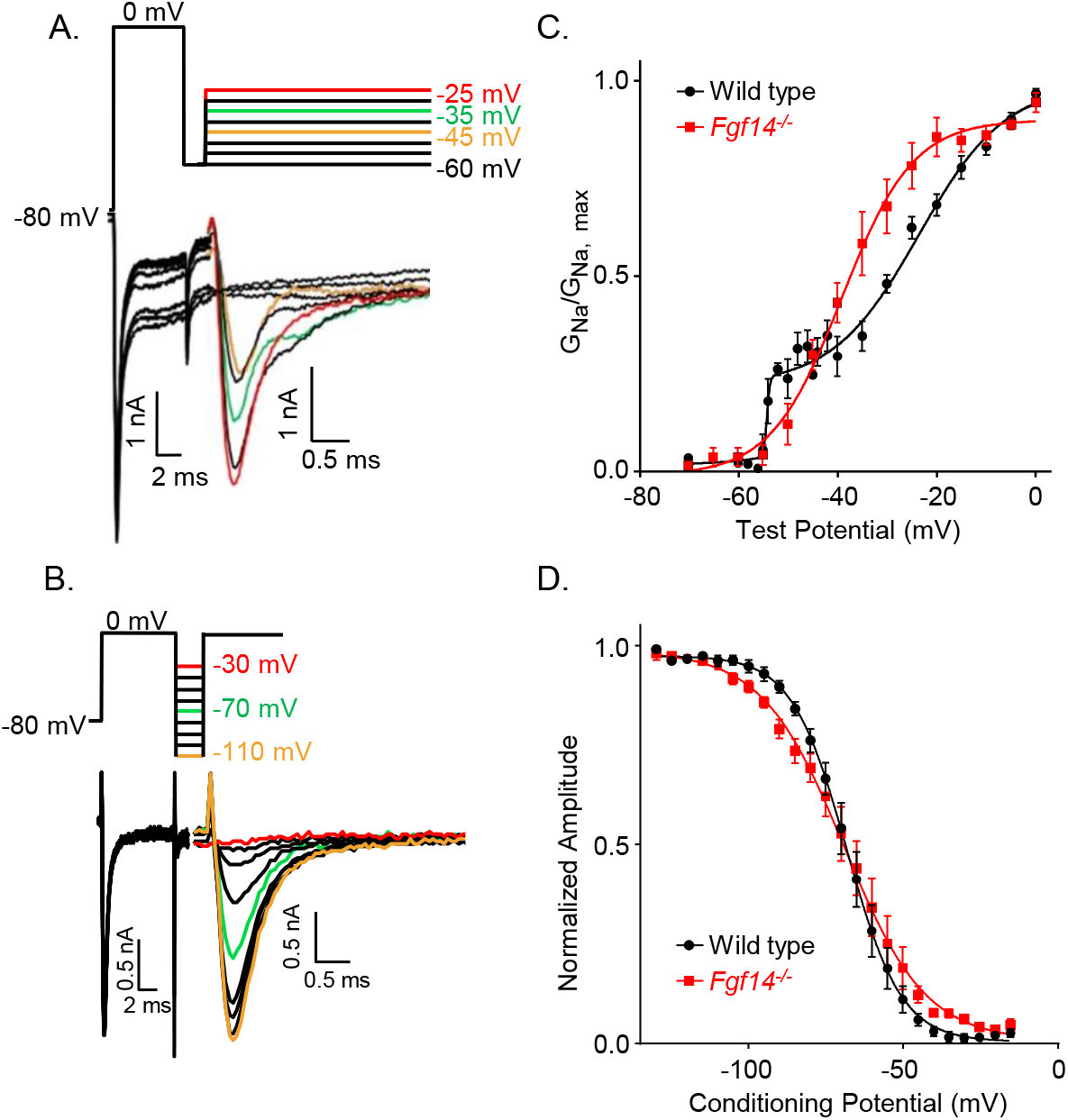
iFGF14 shifts the voltage dependence of Nav current activation in hippocampal CA1 pyramidal neurons. Whole-cell voltage-clamp recordings of Nav currents were obtained from WT and *Fgf14^-/^*^-^ CA1 pyramidal neurons in acute hippocampal slices, prepared from 4-6 week old animals. As described in **Materials and Methods**, voltage-clamp protocols were developed to allow reliable measurements of Nav currents and determination of the voltage dependences of Nav current activation and steady-state inactivation; representative recordings are shown in (**A**) and (**B**), and the voltage-clamp paradigms are illustrated (in the corresponding colors) above the current records. Note that the persistent Nav currents were digitally subtracted and only the inactivating (transient) Nav currents are shown. (**A**) To examine Nav current activation, a 5 ms voltage step to 0 mV from a holding potential of -80 mV was delvered first to inactivate the Nav currents in the distal neurites. Each cell was then hyperpolarized briefly (3 ms) to -60 mV, to allow recovery of Nav channels in the soma and proximal neurites, and subsequently depolarized to various test potentials (-55 mV to 0 mV in 5 mV increments). The peak transient Nav currents evoked at each test potential were measured in each cell, from records such as those in (**A**), conductances were calculated and normalized to the current evoked (in the same cell) at the most depolarized test potential (0 mV). (**C**) Mean ± SEM normalized peak transient Nav current conductances in WT (●; n = 12) and *Fgf14^-/^*^-^ (▪; n = 8) hippocampal CA1 pyramidal neurons were then plotted as a function of the test potential and fitted using Boltzmann equations (see: **Materials and Methods**). The voltage-dependence of activation of the peak transient Nav conductance in WT (●) hippocampal CA1 pyramidal neurons was best fitted by the sum of two Boltzmanns, whereas the data in *Fgf14^-/-^* (▪) hippocampal CA1 pyramidal neurons were best fitted by a single Boltzmann (see text). (**B**) To examine steady-state inactivation of the Nav currents, a 5 ms voltage step to 0 mV was presented to inactivate the Nav currents. Each cell was subsequently hyperpolarized briefly (3 ms) to various conditioning potentials to allow recovery and, finally, depolarized to 0 mV for 5 ms. (**D**) To quantify the voltage-dependences of Nav current inactivation, the peak transient Nav currents evoked at 0 mV from various conditioning voltages (-110 mV to -30 mV) were measured and normalized to the current evoked (in the same cell) from the most hyperpolarized conditioning potential (-110 mV). (**D**) Mean ± SEM normalized peak transient Nav current amplitudes in WT (●; n = 12) and *Fgf14^-/^*^-^ (▪; n = 8) hippocampal CA1 pyramidal neurons were then plotted as a function of the conditioning potential and fitted using the Boltzmann equation (see: **Materials and Methods**). As is evident, the voltage-dependences of steady-state inactivation (**D**) of the Nav currents in *Fgf14^-/-^* (▪) and WT (●) hippocampal CA1 pyramidal neurons are quite similar (see text).

To determine the voltage-dependences of steady-state inactivation of I_NaT_ in WT and *Fgf14^-/-^* CA1 pyramidal neurons, peak Nav currents evoked at 0 mV from various conditioning voltages (**Figure 4B**), were measured and normalized to the amplitude of the peak current evoked from -130 mV (in the same cell). Mean (± SEM) normalized peak Nav currents were then plotted as a function of the conditioning voltage and fitted (**Figure 4D**). The I_NaT_ steady-state inactivation data for both *Fgf14^-/-^* (n = 8) and WT (n = 6) CA1 pyramidal neurons were well described by single Boltzmann functions with similar V_1/2_ = -69.0 ± 1.0 mV and slope = 13.0 ± 1.0 and V_1/2_ = -68.0 ± 0.6 mV and slope = 9.0 ± 0.6 values for *Fgf14^-/-^* and WT, respectively, hippocampal CA1 pyramidal neurons.

### AIS distributions of Ankyrin G and Na1.6 in CA1 pyramidal neurons unaffected by loss of iFGF14

Immunohistochemical analyses of the AIS of mature cerebellar Purkinje neurons *in situ* revealed no measurable effects of the loss of iFGF14 on anti-Ankyrin G or anti-Nav α subunit labelling intensity or distribution (Bosch et al., 2015). It has previously been reported, however, that the acute knockdown of *Fgf14* in dissociated hippocampal neurons in culture disrupts the AIS localization of Nav channels (Pablo et al., 2016). To determine if these seemingly disparate observations reflected differences in neuronal cell types or experimental protocols, we completed experiments to determine directly if iFGF14 affects AIS organization in mature CA1 pyramidal neurons *in situ*. In these experiments, hippocampal sections prepared from adult WT an *Fgf14^-/-^*mice were stained with anti-Ankyrin G and anti-Nav1.6 specific antibodies, and labeling intensities in confocal line scans were determined (see: **Materials and Method**). These experiments revealed robust anti-Ankyrin G and anti-Nav1.6 immunostaining at the AIS of adult WT and *Fgf14^-/-^* hippocampal CA1 pyramidal neurons (**Figure 5A** and **5B**). Quantification of the normalized distributions of both anti-Ankyrin G (**Figure 5C**) and anti-Nav1.6 (**Figure 5D**) immunolabeling along the AIS are not different in WT and *Fgf14^-/-^* CA1 pyramidal neurons. The loss of iFGF14, therefore, does *not* alter the distribution of Nav1.6 at the AIS of adult hippocampal CA1 pyramidal neurons (see: **Discussion**).

**Figure 5.**
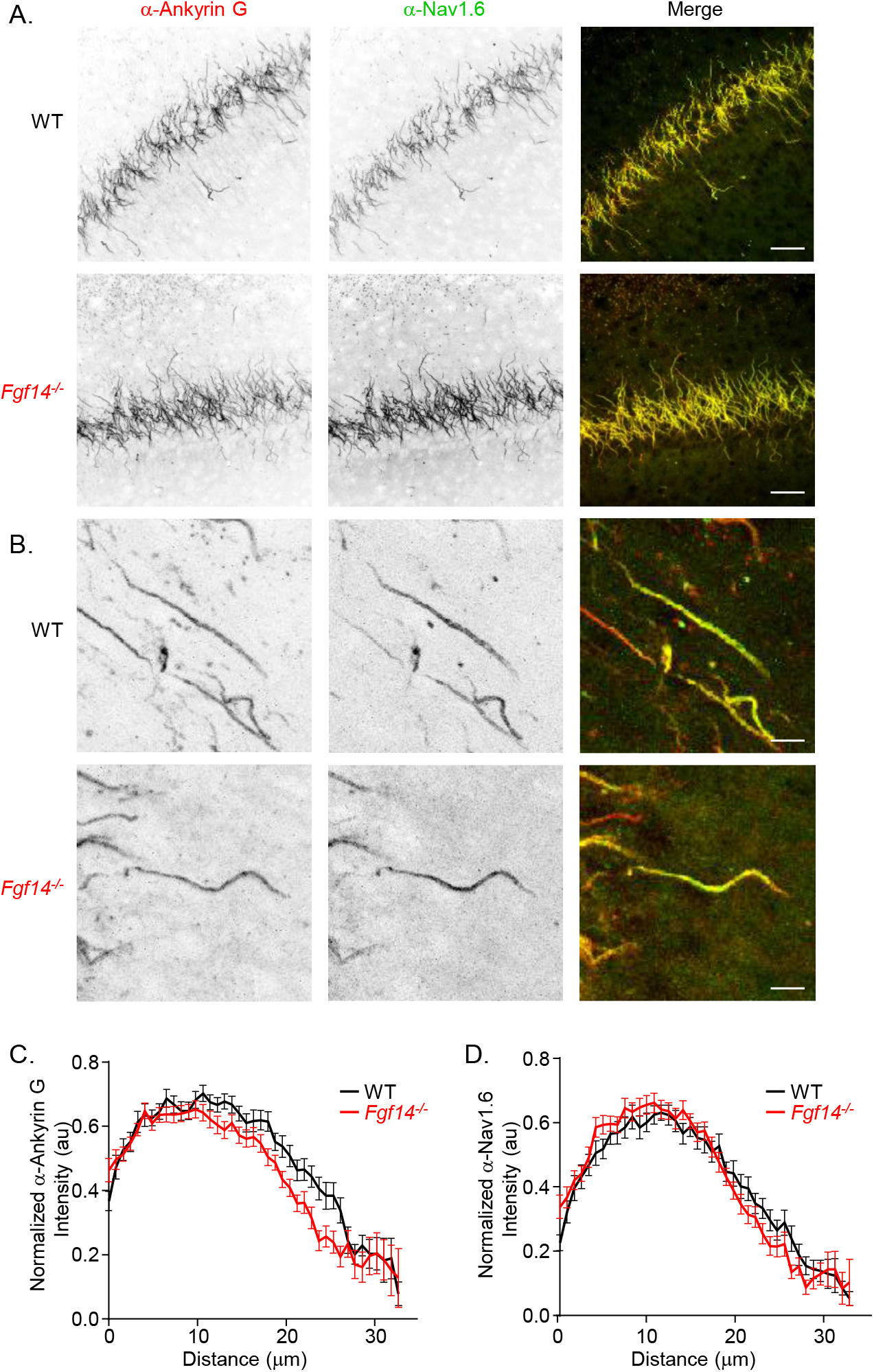
The distributions of anti-Ankyrin G and anti-Nav1.6 labeling along the AIS of WT and *Fgf14^-/-^*hippocampal CA1 neurons are indistinguishable. (**A**,**B**) Representative images of WT and *Fgf14^-/-^* hippocampal CA1 pyramidal neurons in horizontal sections cut from the brains of adult animals and probed with anti-Ankyrin G (red) and anti-Nav1.6 (green) specific monoclonal antibodies, as described in **Materials and Methods**; scale bars are 25 μM in (**A**) and 5 μM in (**B**). (**C**,**D**) The mean ± SEM normalized distributions of anti-Ankyrin G (**C**) and anti-Nav1.6 (**D**) immunofluorescence labeling intensities are indistinguishable along the AIS of *Fgf14^-/-^* (▪; n = 40) and WT (●; n = 32) hippocampal CA1 pyramidal neurons.

### Evoked repetitive firing rates are also *higher* in *Fgf14^-/-^*, than in WT, cortical pyramidal neurons

Additional experiments were undertaken to determine if the loss of iFGF14 increased or decreased the excitability of cortical pyramidal neurons. Whole-cell current-clamp recordings were obtained from visually identified layer 5 pyramidal neurons in acute coronal slices prepared from adult (6-8 week) WT and *Fgf14^-/-^* primary visual cortex (as described in **Materials and Methods**). As illustrated in the representative recordings shown in in **Figure 6A**, WT layer 5 cortical pyramidal neurons fired repetitively in response to prolonged (500 ms) depolarizing current injections, and the rates of repetitive firing increased as a function of the amplitude of the current injection (**Figure 6A**, left panel). Prolonged depolarizing current injections also elicited repetitive firing in *Fgf14^-/-^* layer 5 cortical pyramidal neurons (**Figure 6A**, right panel) and, as in WT cells, the firing rates varied with the amplitude of the injected current. Similar results were obtained in recordings from additional WT and *Fgf14^-/-^* cortical pyramidal neurons, and, similar to the findings in hippocampal CA1 pyramidal neurons (**Figure 2B**), the mean ± SEM input-output curves (numbers of spikes evoked plotted versus injected current amplitude) for WT (n = 16) and *Fgf14^-/-^* (n = 13) cortical pyramidal neurons (**Figure 6B**) are distinct (*P* = 2 x 10^-5^; two-way ANOVA). The mean ± SEM repetitive firing rates determined in layer 5 visual cortical pyramidal neurons (**Figure 6B**) are similar to those measured in CA1 hippocampal pyramidal neurons (**Figure 2B**). Spike frequency adaptation was not found to be different in WT and *Fgf14^-/-^* layer 5 visual cortical pyramidal neurons (**Figure 6A**). Mean ± SEM ratios of firing frequency measured at the end (last 100 ms) and at the beginning (first 100 ms) of 400 pA current injections in *Fgf14^-/-^* (0.65 ± .06, n = 9) and WT (0.52 ± 0.06, n = 12) layer 5 visual cortical pyramidal neurons were *not* significantly different (*P* = 0.13).

**Figure 6.**
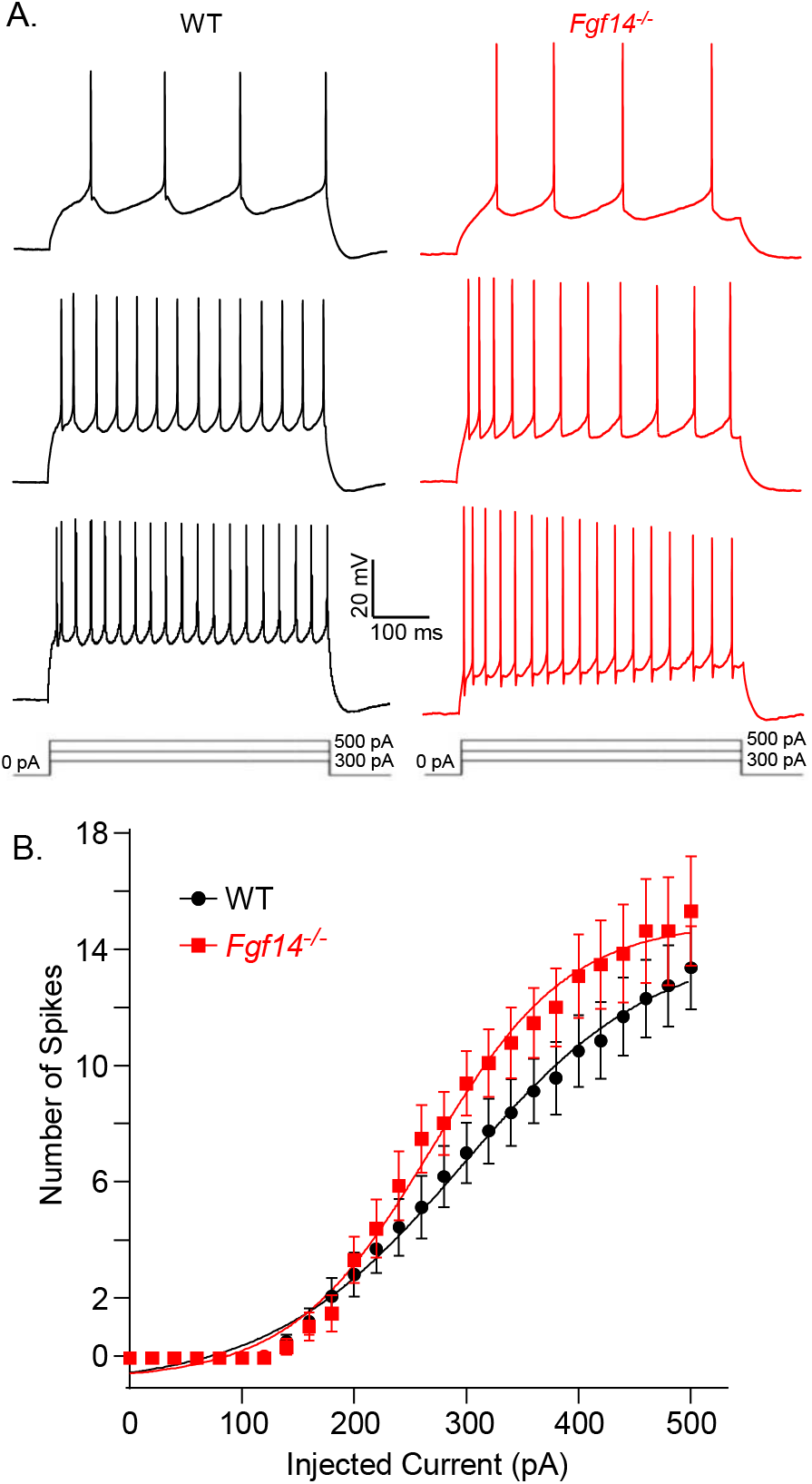
Loss of iFGF14 increases repetitive firing rates in adult layer 5 visual cortical pyramidal neurons. Repetitive firing was evoked in layer 5 visual cortical pyramidal neurons in acute slices prepared from adult (6-8 week) WT and *Fgf14^-/-^* animals, in response to prolonged (500 ms) depolarizing current injections of varying amplitudes (see: **Materials and Methods**). Representative voltage recordings are shown in (**A**) with the injected current amplitudes illustrated below the voltage records. (**B**) Mean ± SEM numbers of action potentials evoked during 500 ms current injections are plotted as a function of the amplitudes of the injected currents. At the higher (>200 pA) injected current amplitudes, the input-output curves (**B**) for *Fgf14^-/-^*(▪; n = 14) and WT (●; n = 16) layer 5 cortical pyramidal neurons are distinct (*P* < 0.01, two-way ANOVA). (**C**) In addition, the mean ± SEM relative amplitude of the last to the first action potential evoked in the train during the 500 pA current injection was lower (\*\**P* < 0.01) in *Fgf14^-/-^*(▪; n = 14), than in WT (●; n = 16), layer 5 visual cortical pyramidal neurons (see text).

### Waveforms of individual action potentials are also altered in *Fgf14^-/-^* cortical pyramidal neurons

As illustrated in the representative records presented in **Figure 7A**, the waveforms of individual action potentials evoked by brief (2 ms) depolarizing current injections in WT and *Fgf14^-/-^* layer 5 visual cortical pyramidal neurons are similar. In contrast with the *depolarizing* shift in the V_thr_ for action potential generation observed in *Fgf14^-/-^*, compared with WT, hippocampal CA1 pyramidal neurons (**Figure 3A** and **3B**), the V_thr_ for action potential generation is more *hyperpolarized* in the *Fgf14^-/-^*, than in the WT, visual cortical pyramidal neuron (**Figure 7A**). Similar results were obtained in recordings from additional WT and *Fgf14^-/-^* layer 5 visual cortical pyramidal neurons, and, as illustrated in **Figure 7B**, the mean ± SEM V_thr_ is markedly lower (*P* = 0.009) in *Fgf14^-/-^*, compared with WT, cortical pyramidal neurons. Similar to hippocampal CA1 pyramidal neurons (**Figure 3C**), however, analyses of action potential durations revealed that the mean ± SEM action potential duration at 50% repolarization is markedly shorter (*P* = 0.025) in *Fgf14^-/-^*, than in WT, cortical pyramidal neurons (**Figure 7C**).

**Figure 7.**
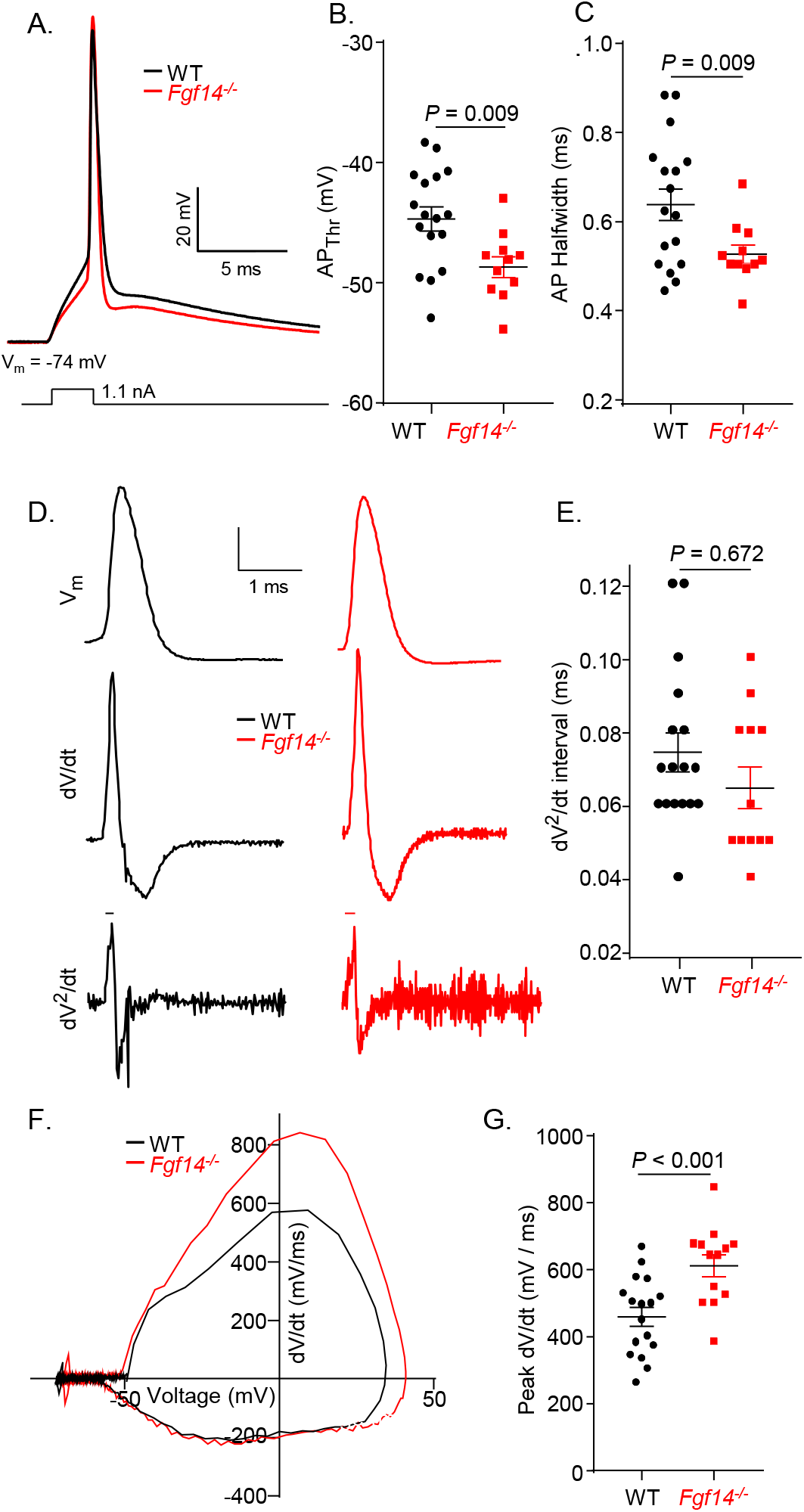
Waveforms of individual action potentials in adult WT and *Fgf14^-/^* layer 5 visual cortical pyramidal neurons are distinct. (**A**) Representative recordings of individual action potentials, evoked in response to brief (2.5 ms) depolarizing current injections (see: **Materials and Methods**), in adult WT (*black*) and *Fgf14^-/-^* (*red*) layer 5 visual cortical pyramidal neurons are shown. The mean ± SEM voltage threshold for action potential generation (AP_Thr_) was more hyperpolarized (**B**) and the mean ± SEM action potential duration at halfwidth (AP_Halfwidth_) was shorter (**C**) in *Fgf14^-/-^*(▪; n = 13), than in WT (●; n = 17), layer 5 cortical pyramidal neurons. (**D**) The representative action potentials in (**A**) are shown on an expanded time scale (*top panels*), and the first (dV/dt; *middle panels*) and second (dV^2^/dt; *lower panels*) derivatives of the action potentials are illustrated in the panels immediately below. The voltage trajectories (dV/dt) of individual action potentials (*middle panels*) were similar in WT and *Fgf14^-/-^*layer 5 cortical pyramidal neurons. In the second (dV^2^/dt) derivatives of the voltage trajectories (*lower panels*), two inflection points were resolved in the rising phases of the action potentials, and the mean ± SEM time interval between these inflection points (**E**) was similar in WT (●; n = 17) and *Fgf14^-/-^*(▪; n = 13) layer 5 cortical pyramidal neurons. (**F**) Construction of phase (dV/dt versus voltage) plots, however, revealed a clear difference in the rising phases of the action potentials in WT and *Fgf14^-/-^*layer 5 cortical pyramidal neurons. As illustrated in (**G**), similar results were obtained in many cells, and the mean ± SEM peak dV/dt value for action potentials evoked in *Fgf14^-/-^* (▪; n = 13) layer 5 cortical pyramidal neurons is higher than in WT (●; n = 17) layer 5 cortical pyramidal neurons.

As illustrated in **Figure 7D** (top traces) and, similar to the findings in CA1 pyramidal neurons (**Figure 3D**), the membrane voltage trajectories during individual action potentials are similar in WT and *Fgf14^-/-^* cortical pyramidal neurons. Distinct from hippocampal CA1 pyramidal neurons, however, the first (dV/dt) (**Figure 7D**, middle traces) and second (dV^2^/dt) (**Figure 7D**, bottom traces) derivative plots of action potential trajectories versus time in WT and *Fgf14^-/-^* cortical pyramidal neurons are similar, i.e., the distinct phases of the action potential upstroke observed in CA1 pyramidal neurons (**Figure 3D**) are not as pronounced in layer 5 cortical pyramidal neurons (**Figure 7D**). Similar to CA1 pyramidal neurons, however, phase plots (**Figure 7F**) revealed that peak dV/dt values are larger (*P* = 0.0007) in *Fgf14^-/-^*, compared with WT, layer 5 visual cortical pyramidal neurons (**Figure 7G**; **Table 1**).

### AIS distribution of Ankyrin G/Na1.6 in cortical pyramidal neurons unaffected by loss of iFGF14

Additional experiments were completed to determine if loss of iFGF14 affects the distribution of Ankyrin G and/or Nav1.6 at the AIS of adult mouse layer 5 visual cortical pyramidal neurons. In these experiments, coronal sections of primary visual cortex prepared from adult WT an *Fgf14^-/-^* mice were stained with anti-Ankyrin G and anti-Nav1.6 specific antibodies, and labeling intensities determined from confocal line scans were compared (see: **Materials and Method**). Similar to hippocampal CA1 pyramidal neurons, the normalized distributions of both anti-Ankyrin G (**Figure 8C**) and anti-Nav1.6 (**Figure 8D**) immunolabeling along the AIS are not different in WT and *Fgf14^-/-^*neurons.

**Figure 8.**
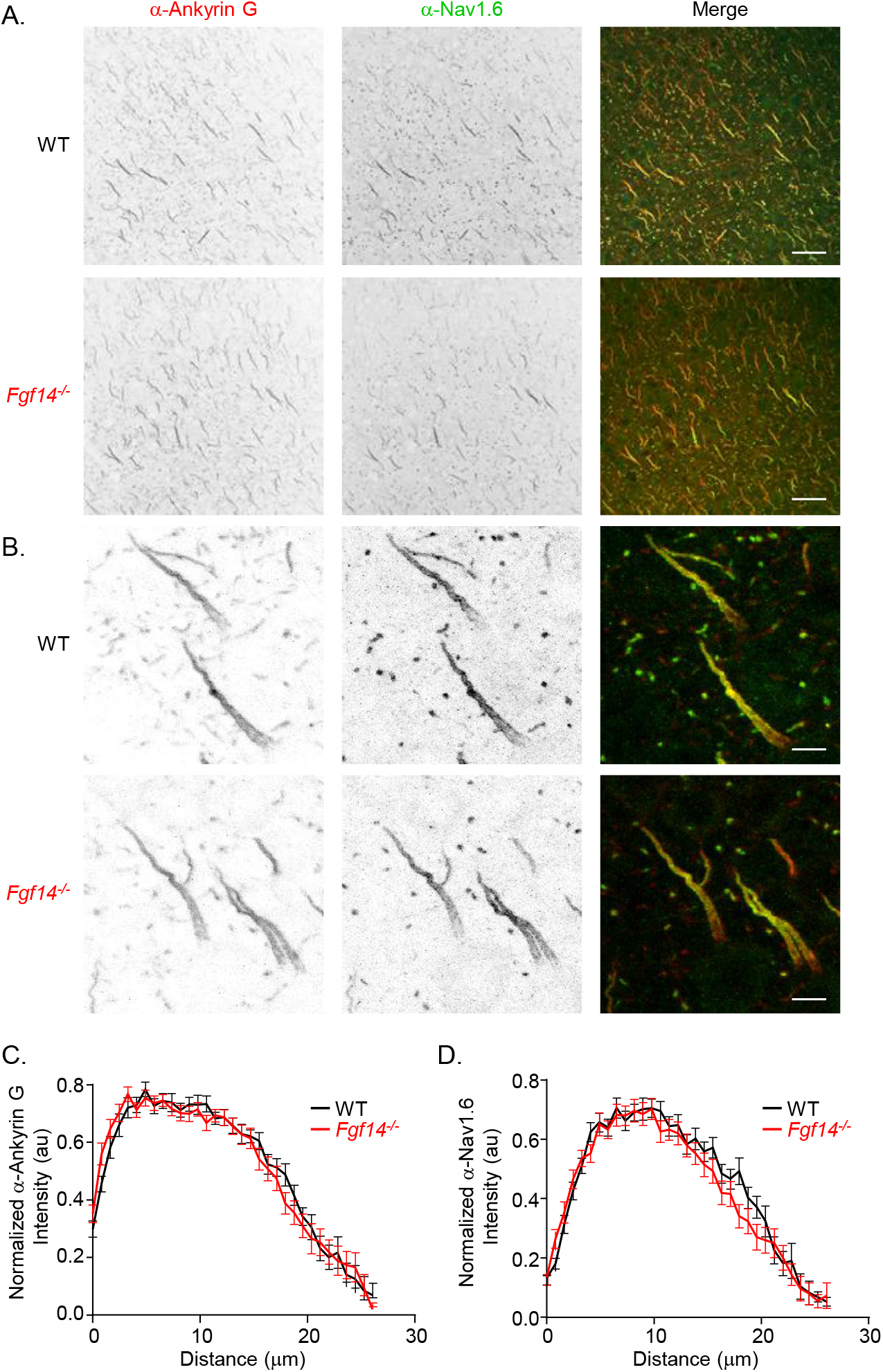
The distributions of anti-Ankyrin G and anti-Nav1.6 labeling along the AIS of WT and *Fgf14^-/-^* layer 5 cortical pyramidal neurons are indistinguishable. (**A**,**B**) Representative images of WT and *Fgf14^-/-^* layer 5 visual cortical pyramidal neurons in horizontal sections cut from the brains of adult animals and probe with anti-Ankyrin G (red) and anti-Nav1.6 (green) specific antibodies as described in **Materials and Methods**; scale bars are 25 μM in (**A**) and 5 μM in (**B**). (**C**,**D**) The mean ± SEM normalized distributions of anti-Ankyrin G (**C**) and anti-Nav1.6 (**D**) immunofluorescence labeling intensities are indistinguishable along the AIS of *Fgf14^-/-^* (▪; n = 25) and WT (●; n = 24) layer 5 cortical pyramidal neurons.

## DISCUSSION

The results of the experiments here demonstrate that evoked repetitive firing rates are *higher* in adult *Fgf14^-/-^*, compared with WT, hippocampal CA1 pyramidal neurons. Marked differences in the waveforms of individual action potentials in adult WT and *Fgf14^-/-^* CA1 pyramidal neurons were also evident: the voltage threshold for action potential generation (AP_thr_) was depolarized in *Fgf14^-/-^*, compared with WT, hippocampal CA1 pyramidal neurons, and the loss of iFGF14 reduced the time delay between the initiation of axonal and somal action potentials. Voltage-clamp experiments revealed that the loss of iFGF14 altered the voltage-dependence of activation, but not inactivation, of I_NaT_ in adult hippocampal CA1 pyramidal neurons. In addition, the immunohistochemical experiments presented show that the loss of iFGF14 does not alter the mean ± SEM normalized distributions of anti-Nav1.6 or anti-ankyrin G immunofluorescence labeling intensities along the AIS of mature hippocampal CA1 pyramidal neurons *in situ*. Taken together, these results demonstrate that, in marked contrast with previous findings reported for iFGF14 knockdown in dissociated neonatal (rat) hippocampal pyramidal neurons in dispersed cell culture (Pablo et al., 2016), the complete loss of iFGF14 throughout development *in vivo does not* measurably affect AIS architecture or the AIS localization of Nav channels in mature hippocampal CA1 pyramidal neurons. Additional experiments and analyses presented here demonstrate that the *in vivo* loss of iFGF14 also resulted in *increased* repetitive firing rates in adult layer 5 visual cortical pyramidal neurons *in situ* without measurably affecting Nav channel AIS localization and distribution.

### Loss of iFGF14 does *not* disrupt the AIS of adult hippocampal pyramidal neurons *in situ*

It has previously been reported that the acute knockdown of iFGF14 in dissociated neonatal (rat) hippocampal pyramidal neurons resulted in reduced Nav current densities and reduced AIS localization of Nav α subunits in spite of preserved AIS architecture, as revealed by the preserved expression of Ankyrin G (Pablo et al., 2016). The results of the immunohistochemical analyses conducted and presented here, however, revealed no significant differences in the normalized distribution of anti-Nav1.6 α subunit labelling of the AIS of *Fgf14^-/-^*, compared with WT, adult (mouse) hippocampal CA1 pyramidal neurons. It seems unlikely that these conflicting results reflect differences in the ages of the neurons studied as the *Fgf14^-/-^* adult hippocampal CA1 pyramidal neurons studied here lack iFGF14 throughout development. In addition, we observed no significant differences in the normalized distribution of anti-Nav1.6 α subunit labelling of the AIS of *Fgf14^-/-^*, compared with WT, adult layer 5 visual cortical pyramidal neurons. It is certainly also possible that these disparate results reflect the fact that mouse hippocampal (and cortical) pyramidal neurons were studied here, whereas Pablo et al. (2016) reported the results of analyses completed on rat hippocampal neurons. A role for cell dissociation is also possible. Additional experiments, designed to compare directly AIS architecture and the impact of the loss of iFGF14 on the AIS localization of Nav channels in mouse and rat hippocampal neurons in dissociated cell culture and in adult mouse hippocampal neurons *in situ*, will be needed to distinguish among these possibilities.

### Distinct effects of iFGF14 in hippocampal/cortical pyramidal, versus cerebellar Purkinje, neurons

The observations here that evoked repetitive firing rates are higher in *Fgf14^-/-^*, compared with WT, adult hippocampal CA1 and layer 5 visual cortical pyramidal neurons and that the voltage-dependence of Nav current activation in CA1 neurons is selectively affected by the loss of *Fgf14* contrast markedly with our previous findings showing that spontaneous repetitive firing rates were significantly reduced in *Fgf14^-/-^*, compared with WT, cerebellar Purkinje neurons (Shakkottai et al., 2019; Bosch et al., 2015), and that this effect was mediated by a hyperpolarizing shift in the voltage-dependence of Nav current inactivation with loss of iFGF14 (Bosch et al., 2015). The motivation for undertaking studies to define the role of iFGF14 in the hippocampus/cortex was that cognitive deficits, like those observed in individuals with SCA27A (Brusse et al., 2006; Misceo et al., 2009), are also evident in *Fgf14^-/-^* animals (Wang et al., 2002; Xiao et al., 2007; Wozniak et al., 2007). We selected hippocampal/cortical pyramidal neurons here for comparison to cerebellar Purkinje neurons because pyramidal cells are functionally similar in that they are, like Purkinje cells (Billard et al., 1993; Gauck and Jaeger, 2000; Ito, 2001), projection neurons. These cell types are, however, distinct in other ways in that cerebellar Purkinje neurons, which express the neurotransmitter GABA (gamma-aminobutyric acid), are inhibitory (Ito et al., 1964; Ito, 2001), whereas hippocampal and cortical pyramidal neurons, which express the neurotransmitter glutamate, are excitatory (Storm-Mathisen, 1981; Jones, 1986). It seems unlikely, however, that the distinct functional effects observed with the loss of iFGF14 in hippocampal/cortical pyramidal neurons, compared with cerebellar Purkinje neurons, simply reflect these differences in neurotransmitter phenotypes as loss of *Fgf14* in cerebellar granule neurons, which are also glutamate-expressing excitatory central neurons (Eccles et al., 1964; Ito, 2001), resulted in a marked *decrease* in evoked repetitive firing rates, attributed to a hyperpolarizing shift in the voltage-dependence of Nav current inactivation (Goldfarb et al., 2007), i.e., cellular phenotypes that are remarkably similar to those we previously described in cerebellar Purkinje neurons (Shakkottai et al., 2009; Bosch et al., 2015).

Studies in heterologous expression systems have demonstrated diverse effects of iFGF co-expression on Nav current amplitudes, as well as on the voltage-dependences of Nav current activation and inactivation (Liu et al., 2001, 2003; Wittmack et al., 2004; Lou et al., 2005; Laezza et al., 2009; Wang et al., 2011b). Comparison of the results obtained in these various studies suggest that the diverse functional effects of each of the iFGFs vary with the specific Nav α subunit and the specific iFGF protein that are co-expressed, as well as with the cellular expression environment, i.e., the effects of an individual iFGF on the expression levels and/or properties of Nav currents encoded by a single Nav α subunit have been shown to be different in different cell types (Wittmack et al., 2004; Lou et al., 2005; Laezza et al., 2007, 2009; Wang et al., 2011b). Given that Nav1.6 is the predominant Na α subunit responsible for the generation of Nav currents at the AIS of central neurons, including hippocampal/cortical pyramidal neurons and cerebellar Purkinje (and granule) neurons (Levin et al., 2006; Royeck et al., 2008; Hu et al., 2009; Dulla and Huguenard, 2009; Katz et al., 2018; Leterrier, 2018; Zybura et al., 2021) and that we are examining the effects of iFGF14 and, more specifically, iFGF14B (**Figure 1**), it seems reasonable to conclude that the distinct functional effects of the targeted deletion of *Fgf14* reflect the different cellular expression environments, i.e., cerebellar Purkinje versus hippocampal/cortical pyramidal neurons. Alternative experimental approaches will need to be developed and employed to determine if the observed functional differences in the modulatory effects of iFGF14 reflect distinct, cell type- (i.e., hippocampal/cortical pyramidal versus cerebellar Purkinje/granule) specific differences in the presence and functioning of other Nav channel accessory subunits, such as accessory Navβ subunits (O’Malley and Isom, 2015; Ransdell et al., 2017) or calmodulin (Ben-Johny et al., 2015) or, possibly, other post-translational modifications (Shao et al., 2009; Onwuli and Beltran-Alvarez, 2016; Pei and Cummins, 2018; Zybura et al., 2021; Marosi et al., 2022) of native Nav channels.

### Physiological and pathophysiological implications

The results presented here demonstrate a physiological role for iFGF14 in the regulation of the intrinsic excitability of adult hippocampal CA1 pyramidal neurons (**Figure 2**) through modulation of the voltage-dependence of Nav channel activation (**Figure 4**). It is unclear, however, whether iFGF14 should be considered an obligatory accessory subunit of Nav channels in hippocampal or cortical pyramidal (and/or in other) neurons and/or if iFGF14-Nav α subunit interactions are dynamically regulated. In addition, it seems reasonable to suggest that, if there are intracellular second messenger pathways (Shao et al., 2009; Onwuli and Beltran-Alvarez, 2016; Zybura et al., 2021; Marosi et al., 2022) that regulate or modulate the interaction(s) between iFGF14 and native neuronal Nav channel α subunits, these could also affect Nav channel activation/availability and, therefore, impact the excitability of hippocampal/cortical pyramidal (and other central) neurons. Delineation of cell signaling pathways that modulate (attenuate or enhance) iFGF14-Nav channel interactions through kinase-mediated and other intracellular second messenger pathways could also provide new strategies to target iFGF-Nav channel interactions and indirectly impact neuronal excitability, potentially in a cell type specific manner.

## Acknowledgements

The authors acknowledge the financial support provided by the National Institute of Neurological Disorders and Stroke of the National Institutes of Health (NIH) (Grant # R01 NS065761 to DMO and JMN; Individual Postdoctoral Fellowship Awards # F32 NS090765 to JLR and # F32 NS065582 to YC). MKB was supported by NIH Institutional Training Grants (T32 GM007200 and T32 HL007275). The monoclonal anti-iFGF14, anti-Ankyrin G and anti-Nav1.6 antibodies were developed by the University of California at Davis/NIH NeuroMab facility, supported by NIH grant U24NS050606, and maintained by the University of California at Davis, and were purchased from Antibodies, Incorporated (Davis CA). The authors declare no competing financial interests.

## Author contributions

YC, JMN, JLR, and DMO conceptualized and planned experiments. Study investigation (experiments and data collection) were performed by YC, JLR, MKB, and RLM. All authors participated in the formal analysis of collected data. The original manuscript draft was written by JLR and JMN. All authors participated in the review and editing of the manuscript.

## FIGURE LEGENDS

**Supplemental Figure 1. Source data for Figure 1.** Full Western blots of fractionated total protein lysates and proteins immunoprecipitated with a mouse monoclonal anti-FGF14 antibody (mAb-α-iFGF14) from the CA1 region of the hippocampus (**A**) and the primary visual cortex (**B**) of adult WT and *Fgf14^-/-^* animals, probed with a rabbit polyclonal iFGF14 antiserum (Rb-α-iFGF14), are shown; the arrowheads indicate the iFGF14 protein (missing in the *Fgf14^-/-^* samples). The regions outlined by the blue boxes indicate the parts of the blots shown in **Figures 1A** and **B**.

**Supplemental Figure 2. Comparisons of the expression levels of the *Fgf11*, *Fgf12,* and *Fgf13* transcripts and of the iFGF12 and iFGF13 proteins in the CA1 region of adult WT and *Fgf14^-/-^*mouse hippocampus.** (**A**) qRT-PCR analyses of *Fgf11*, *Fgf12, and Fgf13* transcripts in the CA1 region of the hippocampus of adult WT (●; n = 6) and *Fgf14^-/-^* (▪, n = 6) mice demonstrate robust expression of *Fgf12* and *Fgf13* in both WT and *Fgf14^-/-^* CA1, whereas *Fgf11* is barely detectable. Further analysis, using isoform specific primers (see: **Supplemental Table 1**), reveal that *Fgf12B* is the predominate *Fgf12* isoform and that *Fgf13A* is the predominate *Fgf13* isoform expressed in the CA1 region of both WT and *Fgf14^-/-^* adult mouse hippocampus and, in addition, that the expression levels of the *Fgf12B* and the *Fgf13A* transcripts in WT and *Fgf14^-/-^* CA1 are quite similar. (**B**, **C**) Western blots of fractionated total protein lysates prepared from the CA1 region of adult WT and *Fgf14^-/-^* mice, probed with a (**B**, top panel) rabbit polyclonal anti-iFGF13 antiserum (Rb-α-iFGF13) or a (**C**, top panel) rabbit polyclonal anti-iFGF12 antibody (Rb-α-iFGF12); the arrowheads indicate the iFGF13 (**B**, top panel) and iFGF12 (**C**, lower panel) proteins. To ensure equal protein loading of all lanes, the membranes were also probed with a mouse monoclonal antibody targeting GAPDH (mAb-α-GAPDH) (**B**, lower panel and **C**, lower panel); the arrowheads indicate the GAPDH protein. The full Western blots are shown in **Supplemental Figure 3**.

**Supplemental Figure 3. Source data for Supplemental Figure 2.** Full Western blots of the fractionated total protein lysates from the CA1 region of the hippocampus of adult WT and *Fgf14^-/-^*animals probed with the Rb-α-iFGF13 (**A**) or Rb-α-iFGF12 (**B**) antibody, are shown; the arrowheads indicate the iFGF13 (**A**, left panel) or iFGF12 (**B**, left panel) proteins. Membranes were also probed with a mAb-α-GAPDH, as shown in the right panels of (**A**) and (**B**). The regions outlined by the blue boxes in all four panels indicate the parts of the blots shown in **Supplemental Figure 2B** and **2C**.

**Supplemental Figure 4. Comparisons of the expression levels of the *Fgf11*, *Fgf12,* and *Fgf13* transcripts and of the iFGF12 and iFGF13 proteins in layer 5 of the primary visual cortex of adult WT and *Fgf14^-/-^* mice.** (**A**) qRT-PCR analyses of *Fgf11*, *Fgf12, and Fgf13* transcript expression levels in layer 5 of the primary visual cortex of adult WT (●; n = 6) and *Fgf14^-/-^* (▪, n = 6) mice demonstrate robust expression of *Fgf12* and *Fgf13* in WT and *Fgf14^-/-^*cortex, whereas *Fgf11* is barely detectable. Further analysis, using isoform specific primers (see: **Supplemental Table 1**), demonstrate that *Fgf12B* is the predominate *Fgf12* isoform and that *Fgf13A* is the predominate *Fgf13* isoform expressed in layer 5 of the visual cortex of both WT and *Fgf14^-/-^* animals and that the expression levels of both *Fgf12B* and *Fgf13A* are similar in the WT and *Fgf14^-/-^* samples. (**B**, **C**) Western blots of fractionated total protein lysates prepared from layer 5 of the primary visual cortex of adult WT and *Fgf14^-/-^* mice, probed with the Rb-α-iFGF13 (**B**, top panel) or Rb-α-iFGF12 (**C**, top panel) antibody; the arrowheads indicate the iFGF13 (**B**, top panel) and iFGF12 (**C**, top panel) proteins. The membranes were also probed with mAb-α-GAPDH (**B**, lower panel and **C**, lower panel), and the arrowheads indicate the GAPDH protein. The full Western blots are presented in **Supplemental Figure 5**.

**Supplemental Figure 5. Source data for Supplemental Figure 4.** Full Western blots of fractionated total protein lysates from the primary visual cortex of adult WT and *Fgf14^-/-^*animals, probed with Rb-α-FGF13 (**A**, left panel) or Rb-α-iFGF12 (**B**, left panel), are shown; the arrowheads indicate the iFGF13 (**A**, left panel) and iFGF12 (**B**, left panel) proteins. Membranes were also probed with mAb-α-GAPDH, as shown in the right panels of (**A**) and (**B**), and the arrows in the right panels indicate GAPDH protein. The regions outlined by the blue boxes in all four panels indicate the parts of the blots shown in **Supplemental Figure 4B** and **4C**.

