## Supplementary material for "Loss of Intracellular Fibroblast Growth Factor 14 (iFGF14) *Increases* the Excitability of Mature Hippocampal and Cortical Pyramidal Neurons": Supp. Figures and Table

**Supplemental Table1.** Primers for Quantitative Real Time PCR

| Gene | Forward Primer | Reverse Primer |
|---|---|---|
| *Fgf14* | 5’ caattccacactgttcaacctcat 3’ | 5’ actccctggatggcaacaac 3’ |
| *Fgf14A* | 5’ ggcaacctggtggatatcttctc 3’ | 5’ ggcaacaacgcgcagtc 3’ |
| *Fgf14B* | 5’ ttttcgcccaaatcaatgtg 3’ | 5’ ggcaacaacgcgcagtc 3’ |
| *Fgf11* | 5’ cctcagctcaaaggcatcgt 3’ | 5’ attcgcctggaggtagaaacc 3’ |
| *Fgf12* | 5’ aagggattgtgacaaggttattcag 3’ | 5’ acacgcagtcctacaggaattaga 3’ |
| *Fgf12A* | 5’ ttcagcaaagtgcgcttctg 3’ | 5’ acacgcagtcctacaggaattaga 3’ |
| *Fgf12B* | 5’ tgaagcgggcccacat 3’ | 5’ acacgcagtcctacaggaattaga 3’ |
| *Fgf13* | 5’ gggtggtggctattcaagga 3’ | 5’ gtatccctcgctgttcattgc 3’ |
| *Fgf13A* | 5’ ccagctgcgacaaaaacaagtta 3’ | 5’ gcctgcagttgcaagtggtag 3’ |
| *FgfVY* | 5’ aaagaaaacacagaacccgaagag 3’ | 5’ gcctgcagttgcaagtggtag 3’ |
| *Fgf13B* | 5’ ggaagtcatattcagagcctcagc 3’ | 5’ gcctgcagttgcaagtggtag 3’ |
| *Hprt* | 5’ tgaatcacgtttgtgtcattagtga 3’ | 5’ ttcaacttgcgctcatcttagg 3’ |
| *Pgk-1* | 5’ gcttctggaaacaaggttaaagct 3’ | 5’ acagtgaggctcggaaagca 3’ |

A. CA1

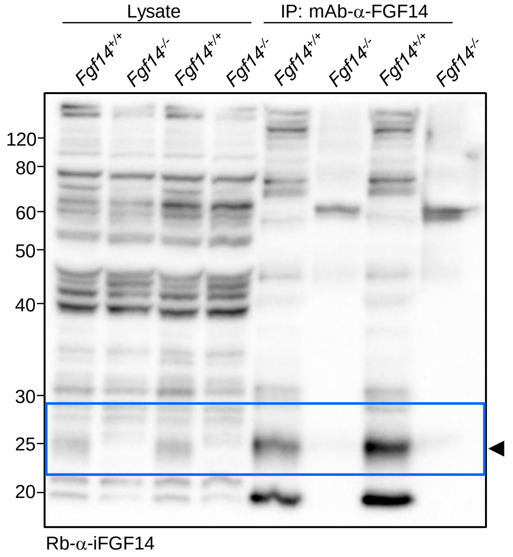


B. Cortex

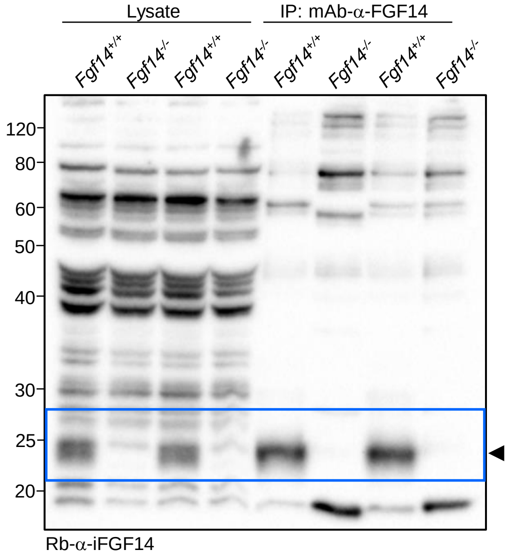


Supplemental Figure 1. Ransdell et al.

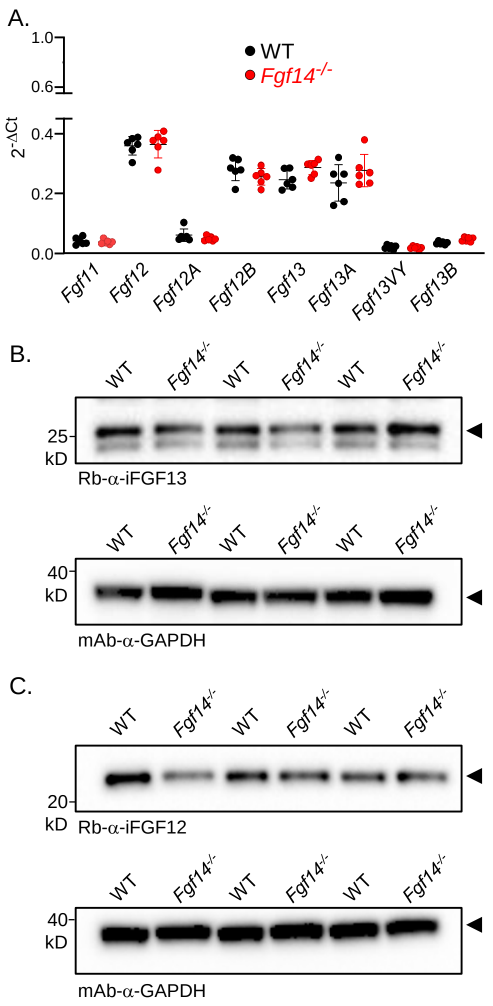


Supplemental Figure 2. Ransdell et al.

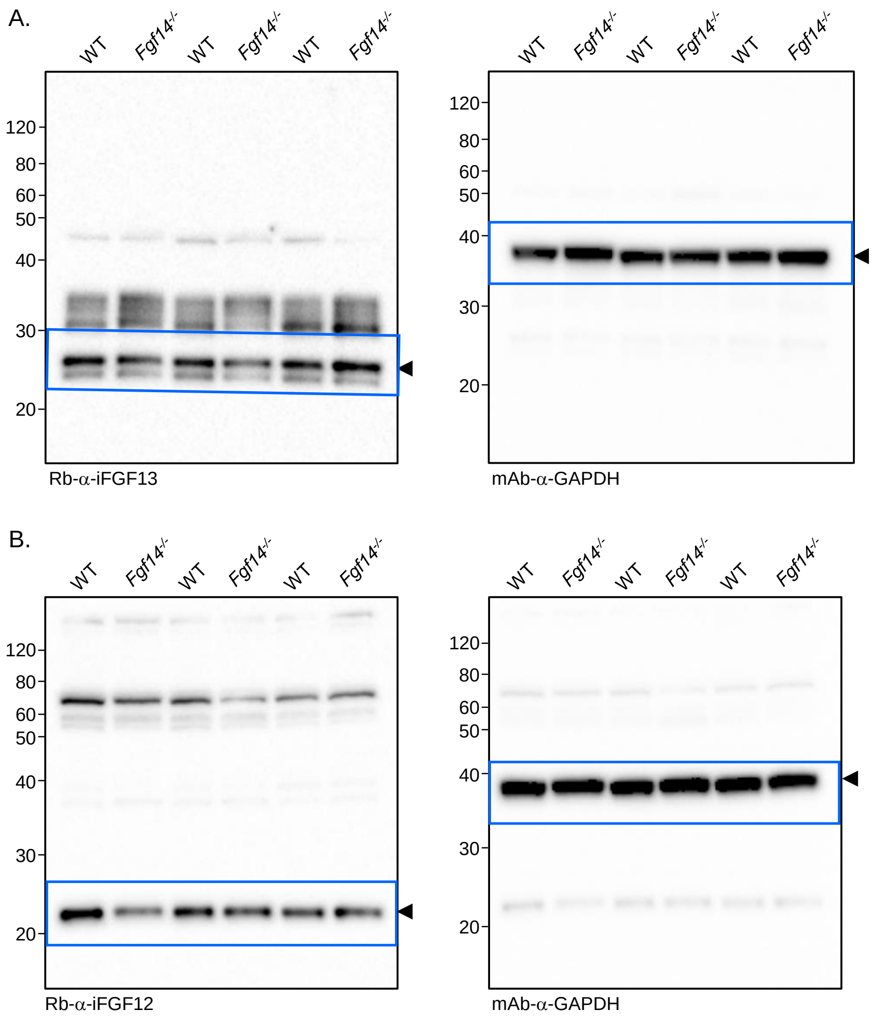


Supplemental Figure 3. Ransdell et al.

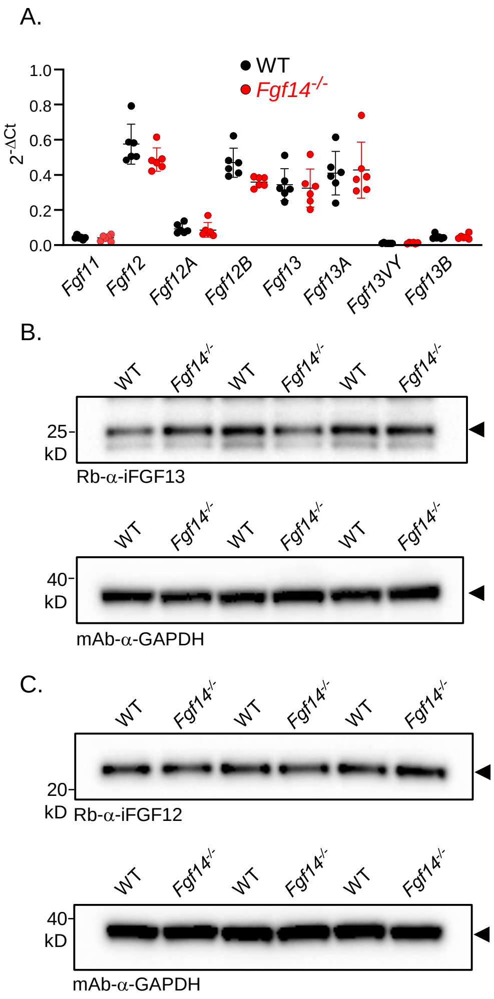


Supplemental Figure 4. Ransdell et al.

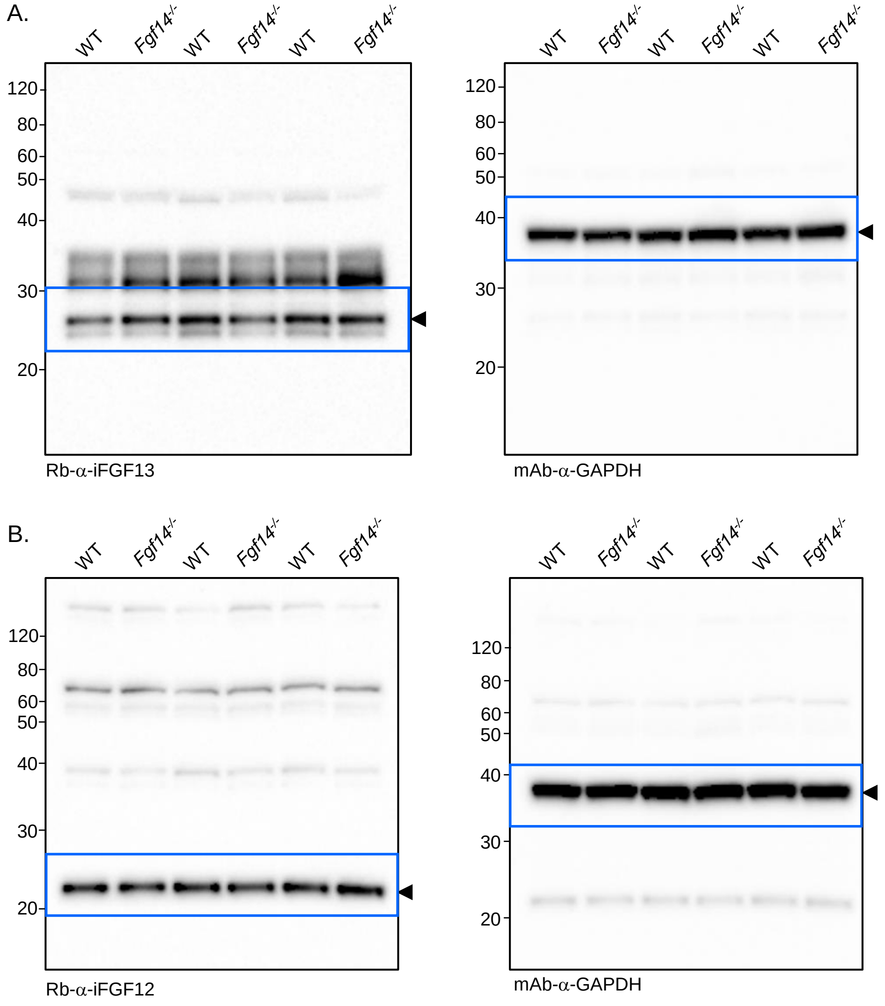


Supplemental Figure 5. Ransdell et al.
